# Patient-specific iPSC models of neural tube defects identify deficiencies in neuroepithelial cell shape regulation and differentiation

**DOI:** 10.1101/2025.05.16.654453

**Authors:** Ioakeim Ampartzidis, Elliott M Thompson, Yashica Gupta, Andrea Krstevski, Nicola Elvassore, Eirini Maniou, Paolo de Coppi, Gabriel L Galea

**Author notes:** Department of Biochemistry, University of Cambridge, Hopkins Building, Tennis Court Road, Cambridge CB2 1QW, UK.

## Abstract

Spina bifida and anencephaly are neural tube defects caused by failure of embryonic neural tube closure. Successful closure requires apicobasal elongation and apical constriction of neuroepithelial cells. These critical behaviours have not been studied in patient-specific models. We characterise a human iPSC-derived neuroepithelial morphogenesis model that is reproducible across parental iPSC lines of diverse origin and reprogramming technologies. Differentiated neuroepithelial cells undergo ROCK-dependent apical constriction. ROCK acts downstream of planar cell polarity/VANGL2 in other species. We find that VANGL2 is up-regulated and phosphorylated during iPSCs-to-neuroepithelial differentiation. The patient-identified *VANGL2-R353C* mutation does not alter VANGL2 expression, but reduces myosin-II phosphorylation and apical constriction relative to congenic controls. Non-congenic comparisons and forward genetic testing are also informative in this reproducible model. We compare lines reprogrammed from amniocytes of two patients with spina bifida, versus two controls. One patient-derived line forms a morphologically normal neuroepithelium, but fails to differentiate into neurons. The second fails to undergo apicobasal elongation, and harbours compound heterozygous mutations in the *MED24* gene implicated in neuroepithelial elongation in mice. Thus, iPSC-derived neuroepithelial modelling reveals mechanistic insights into conserved cell behaviours, including apical constriction, apicobasal elongation and neural differentiation, links genetically-impaired apical constriction to human disease, and establishes patient-specific models.

**Impact statement:** Neural tube defects are common malformations caused by multi-gene/environment interactions. The highly reproducible iPSC-to-neuroepithelium differentiation protocol characterised here provides mechanistic insight into fundamental cell behaviours and enables patient-specific disease modelling.

## Introduction

Neural tube defects (NTDs), such as spina bifida and anencephaly, affect approximately 1:1,000 births globally^1^. These severe neurodevelopmental malformations are caused by abnormal morphogenesis of the neural tube during embryonic development. The most severe NTDs result from failure of the physical process of neural tube closure, which in humans is completed by approximately 30 days gestation^2,3^. Its completion during largely inaccessible stages of pregnancy has limited our ability to study neural tube closure in humans. This challenge is compounded by the multigenic causation of NTDs, as well as interactions of both maternal and embryonic genetics with environmental factors^4–6^. Consequently, few patients receive an interpretable genetic diagnosis.

A few patient-identified mutations have been functionally validated in transgenic animal models^5,7–9^. Mice and other vertebrates have been invaluable models for understanding the process of neural tube closure. Both mice and humans close their open neural tube regions – called neuropores – through progressive elevation of neural folds which meet at the dorsal midline and zipper along the body axis^2,10^. Progression of closure requires the action of different cell types, particularly the surface ectoderm (future epidermis) and neuroepithelium (future brain and spinal cord)^3^. These cells coordinate multiple behaviours within and between each other. Neuroepithelial cells thicken along their apicobasal axis as they differentiate into a pseudostratified epithelium which undergoes interkinetic nuclear migration synchronised with cell cycle progression, superimposed on active apical constriction^11–17^.

Neuroepithelial apical constriction is an evolutionarily conserved force-generating mechanism which contributes to neural tube closure in chordates including amphibians, birds and mammals^3^. It requires targeted localisation and phosphorylation of non-muscle myosin II in the cell’s apical domain. Myosin binds filamentous (F-)actin, cross-linking filaments and generating contractile mechanical forces that pull actin filaments together through power strokes^18^. Neuroepithelial apical enrichment of F-actin and phosphorylation of myosin both require Rho-associated kinase (ROCK)^19^, which is in turn apically enriched at mediolateral cell borders at least in part by signalling through the non-canonical wingless homology (Wnt)/planar cell polarity (PCP) pathway^20–22^. PCP components promote mediolaterally oriented neuroepithelial apical constriction in *Xenopus* frogs^23,24^, chickens^22^ and mice^15,20^, but this pathway has been difficult to study in human epithelial models due to lack of quantifiable readouts relevant to its functions *in vivo*.

PCP was originally described in *Drosophila* as a regulator of polarised cell features such as hair cell bristle orientation^25–27^. Pathway components were later shown to be conserved in vertebrates and contribute to neural tube closure^28–31^. The core PCP pathway involves signalling through asymmetric receptor complexes on adjacent cells, pairing van Gogh-like (VANGL) and Cadherin EGF LAG seven-pass G-type receptor (CELSR) proteins on one cell, with frizzled (FZD)/CELSR complexes on the adjacent cell^32^. Both VANGL and FZD branches of this pathway are involved in neural tube closure^33,34^. VANGL2 can be activated through phosphorylation following binding of PCP Wnt ligands^35,36^.

PCP signalling drives two functionally distinct processes in neuroepithelial cells essential for mammalian NT closure: early convergent extension and later apical constriction. Global genetic deletion of *Vangl2* precludes medial convergence and rostro-caudal elongation necessary to initiate neural tube closure in mouse embryos^37^, although this effect may be partly mediated by the adjacent mesoderm^38^. At later developmental stages, however, VANGL2 promotes cytoskeletal reorganisation involving F-actin planar polarisation and phosphorylation of non-muscle myosin IIB, likely by ROCK^20,39–41^. We previously reported that VANGL2 is not cell-autonomously required for apical constriction of mouse neuroepithelial cells^20^.

Mutations in PCP genes including *VANGL2*^42–47^ have been identified in patients who have NTDs and damaging variants in actomyosin-related cytoskeletal regulation genes are enriched in individuals with spina bifida^48^. However, it has been challenging to study the regulation of apical constriction in human neuroepithelial cell models. A previous study described elegant modelling of neural tube closure by differentiation of induced pluripotent stem cells (iPSCs) on micropatterned surfaces^49^. That model’s formation of neuroepithelium enveloped within a larger cyst limits access to its apical surface for functional measurements, such as inference of mechanical tension through laser ablation assays. Similarly, previous studies have demonstrated that disruption of actomyosin regulation dilates apical surfaces on the inside of human iPSC-derived neuroepithelial spheres^50,51^. Several models have also been described which enable studies into later stages of neuroepithelial function as they differentiate into neural progenitors and then neurons^52–55^. While these neurogenesis models provide insights into later differentiation, they do not capture cell behaviours crucial for neural tube closure.

We recently characterised a minimal model of neuroepithelial morphogenesis in which a monolayer of cuboidal iPSCs apicobasally elongate into a complex, pseudostratified neuroepithelium with a planar morphology^17^. Importantly, this model presents readily accessible neuroepithelial apical surfaces which are directly comparable in size with those of mouse embryos^17^. Moreover, the same protocol can be extended to differentiate neuroepithelial cells into cortical neurons^56^. This enables sequential assessment of neuroepithelial functions relevant to generating mechanical deformations which close the neural tube and their subsequent differentiation which normally occurs after completion of closure. Here, we show that neuroepithelial development in this model is reproducible, even when comparing non-congenic iPSC lines. We validate use of the model to study PCP components including ROCK and VANGL2, finding that a *VANGL2* point mutation previously identified in a patient^46^ diminishes neuroepithelial apical constriction. Extension of the model to two iPSC lines derived from individuals who have spina bifida identifies line-specific deficiencies, including apicobasal elongation and neural differentiation, potentially relevant to their malformation. Strikingly, all three NTD-relevant lines – *VANGL2* point mutation and two patient-derived lines – present failures of distinct neuroepithelial functions, recapitulating the heterogeneity of these conditions.

## Results

### Quantitative, reproducible platform of iPSC-to-neuroepithelial differentiation

Intra-line variability often limits the use of human iPSCs to model multigenic diseases, such as NTDs, in which congenic controls are often impractical. In contrast, we find that neuroepithelial differentiation through dual SMAD inhibition (Figure 1a)^17,56^ is highly reproducible between different iPSC lines. We illustrate this by using three previously reported iPSC lines from healthy individuals, selected to represent different origins and reprogramming methods (Figure 1b). For each of the three iPSC lines, we analysed three independently differentiated plates, each representing a separate biological replicate. Line H0193b was reprogrammed from male (46XY) amniocytes through microfluidic delivery of modified mRNA^57,58^. SFC086-03-01 was reprogrammed from adult female (46XX) fibroblasts using non-integrating Sendai virus^17,59^. 802-30F was from endothelial cells of a Hispanic female (46XX) reprogrammed through StemRNA^60^. Despite their varied origins, all three undergo equivalent, temporally predictable changes in epithelial morphology during neuroepithelial differentiation (Figure 1a-g).

**Figure 1:**
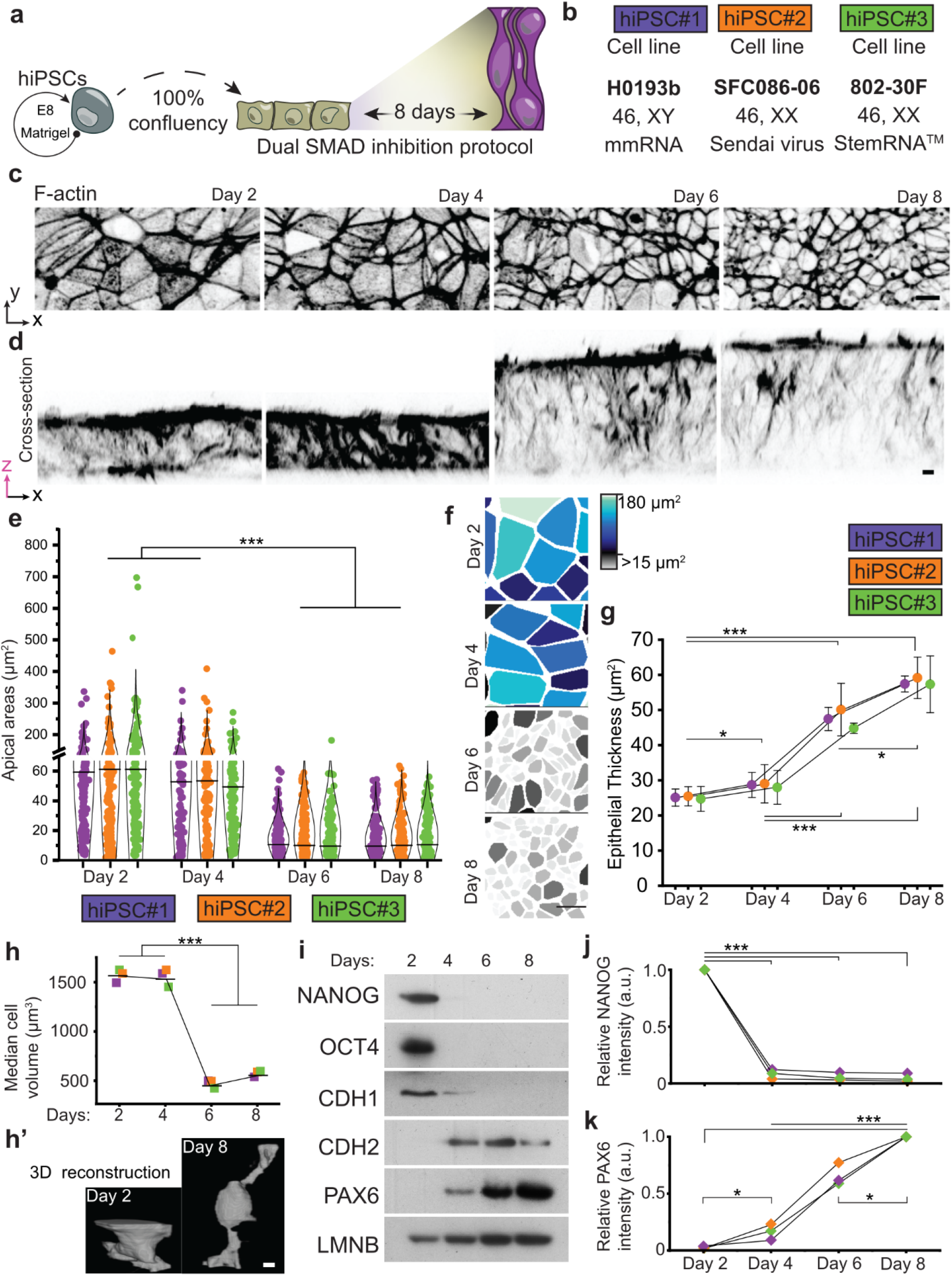
Reproducible morphological and molecular evolution of iPSC differentiation into neuroepithelial cells. iPSC1 is HO193b, iPSC2 is SFC086-03-01 and iPSC3 is 802-30F. **a,** Schematic representation of the neuroepithelial induction protocol. Sub-confluent human iPSCs (in grey) are passaged and seeded at full confluency from the start of differentiation and cultured for eight days, until reaching neuroepithelial maturity. Further passage at this stage forms neural rosettes and allows further differentiation into post-mitotic neurons^56^. **b,** Summary of control lines used to validate reproducibility of differentiation into neuroepithelial cells. **c,d,** Representative high-resolution confocal top-view (**c**) and cross-section (**d**) images of day 2-8 cultures fluorescently labelled to show F-actin with inverted gray LUT. P values from one-way ANOVA with post-hoc Bonferroni. **e,** Dot plot representing cell’s apical area across timepoints of neuroepithelial differentiation, in three iPSC lines. Black bars represent the median apical area across three independent culture plates of each line. **f,** Heatmaps of hiPSC2 representing the apical area size changes during differentiation, in a contiguous area of 1,200 μm^2^ per timepoint. **g,** Quantification of epithelial thickness across neuroepithelial differentiation of control cell lines. Three culture replicates were measured for each time point and cell line. **h,** Dot plot representing the cell volume decrease across timepoints of differentiation with representative 3D volumetric images at day 2 and day 8 (**h’**). **i,** Representative western blot panel of NANOG, OCT4, CDH1, CDH2, PAX6, and LMNB protein levels across timepoints of neuroepithelial differentiation, in the iPSC2 line. **k,j,** Dot plots representing normalised band intensity of NANOG (j), and PAX6 (k), across differentiation for three control cell lines. NANOG expression is normalised to day 2 per line, whereas PAX6 is normalised to day 8 as it is not detected at day 2. P values from one-way ANOVA (**e,g**) or Wilcoxon paired test (**j,k**) with Bonferroni correction (***: p ≤ 0.001, *: p ≤ 0.005). Scale bar = 10 μm.

These cells begin their differentiation as an apicobasally polarised cuboidal monolayer approximately 25 µm thick, with large apical areas approximately 60 µm^2^ (‘day 2’, Figure 1c-g). During differentiation, they double their apicobasal length into a pseudostratified epithelium^17^ approximately 55 µm thick, with small apical surfaces around 10 µm^2^ (‘day 8’, Figure 1c-g, Supplementary Figure 1a-c). Apicobasal elongation increases total epithelial volume, accommodating ongoing cell division^17^, yet the volume of individual cells decreases from 1,550 µm^3^ at the start of differentiation to 500 µm^3^ at the end (Figure 1h). This reduction in total cell volume is partly due to a halving of nuclear volume (Supplementary Figure 1d-e).

Neuroepithelial apical size and apicobasal thickness – properties whose dynamic control enables neural tube closure *in vivo*^20,22,61,62^ – are indistinguishable between non-congenic iPSC lines at each stage of differentiation analysed (Figure 1c-g). Equivalent neuroepithelial thickness and apical areas are also obtained in cultures differentiated by different investigators (Supplementary Figure 2a-b). All iPSC lines show a non-linear morphological evolution during differentiation: apical area and apicobasal thickness remain similar between days 2 and 4 of differentiation, but then change rapidly from day 5 (Supplementary Figure 1a-c) until the end of differentiation on day 8 (Figure 1e-g). In contrast, their expression of pluripotency markers such as NANOG and OCT4 are rapidly lost between days 2 and 4 of differentiation (Figure 1i,j)^17^. The cadherin CDH1 is similarly down-regulated early during differentiation, switching to expression of the neuroepithelial CDH2 from day 4 of differentiation (Figure 1i), before morphological changes occur. The neuroepithelial marker PAX6 is detectably expressed from day 4 and its expression increases between day 4 and 8 (Figure 1i,k).

### Cadherin switching precedes morphological changes during iPSC to neuroepithelium differentiation

We next tested whether the expression of differentiation markers changes synchronously with the onset of apical constriction using cross-correlation analysis. CDH1 down-regulation and the increase in CDH2:CDH1 ratio precede apical area constriction in all three lines tested (Supplementary Figure 2c). In contrast, PAX6 up-regulation is synchronous with apical constriction in all three lines (Supplementary Figure 2c). However, these analyses are based on four timepoints from our western blotting analysis, limiting comparison of correlations.

To explore gene/morphology correlations more globally, we performed bulk RNA sequencing across five pooled samples per day of differentiation, using iPSC3 cells. Cultures were lysed for RNA analysis in parallel with cultures fixed for concomitant immunofluorescence staining and apical area analysis (Figure 2a). We first validated that the profile of five markers examined via western blot (Figure 1i) follows a similar pattern of RNA expression (Figure 2b). All five show RNA expression changes consistent with protein level changes previously observed.

**Figure 2:**
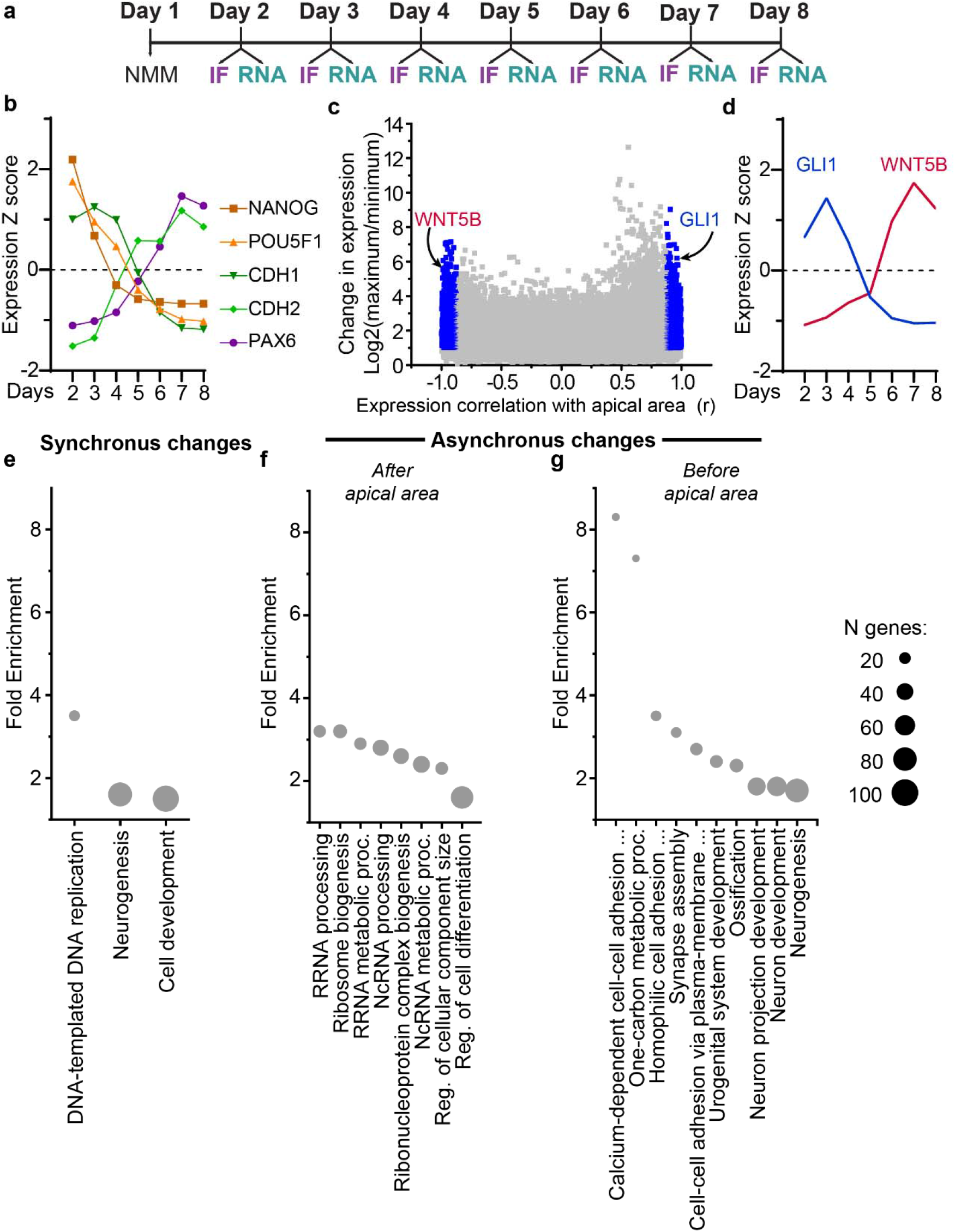
Cross-correlation analysis of gene expression with apical area. a, Schematic representation of gene/morphology correlation protocol. For each day of the protocol five cultures were lysed to extract RNA and one was fixed for immunofluorescence (IF) staining and apical area analysis. NMM indicates cells in neural maintenance medium, such that the first timepoint analysed here (day 2 of differentiation) represents 24 hrs in neural induction medium. **b,** Expression profiles of NANOG, POU5F1, CDH1, CDH2, and PAX6. **c,** Volcano-style plot illustrating the correlation coefficient versus apical area, and change in expression of all genes detected (individual points). Blue points have a fold change >2 and FDR-adjusted correlation p-value <0.05. **d,** Expression profiles of GLI1 and WNT5B. **e, f,g,** Gene ontology enrichment of genes whose expression changes are most strongly correlated with apical area when (**e**) synchronised, (**f**) when offset by +1 day (gene change later than apical area change), or (**g**) when offset by -1 day (genes change earlier than apical area change).

The expression of 3,013 genes changes by ≥2-fold during differentiation and is significantly correlated with apical area (Figure 2c, absolute Pearson r value corresponding to FDR-corrected p < 0.05 for the area vs expression regression slope). This exploratory analysis is not intended to detect genes differentially expressed between days of differentiation. Rather it identifies transcripts whose change in expression is significantly correlated with the non-linear temporal change in apical area. Genes positively correlated with apical area decrease in expression over differentiation (e.g. *GLI1*) whereas those negatively correlated with apical area increase over time (e.g. *WNT5B*, Figure 2d). Overall, genes whose expression changes in synchrony with apical area are enriched in *cell replication* and *neurogenesis*-related gene ontology (GO) terms (Figure 2e). *Proliferation*-related terms are enriched in down-regulated genes, whereas *neurogenesis* genes are enriched in up-regulated genes.

We next sought to identify changes in transcription that occur after the reduction in apical area. These are enriched for *rRNA processing* (down-regulated genes) and *regulation of cell differentiation* (up-regulated genes) GO terms (Figure 2f). Genes whose expression changes before apical area decreases may be responsible for preparing cells to apically constrict. They show the highest fold enrichment for *calcium-dependent cell-cell adhesion* (up-regulated genes, Figure 2g), consistent with our observation of cadherin switching before apical constriction. The second most-enriched GO term is *one-carbon metabolism* process (Figure 2g), related to the folate one-carbon metabolic pathway known to influence neural tube closure. This GO term is not enriched in either the up-regulated or down-regulated genes when assessed in isolation but relevant genes such as *MTHFR* and *DHFR* are down-regulated during differentiation (Supplementary Table 1).

Taken together, these analyses suggest that the expression of neural differentiation markers parallels neuroepithelial morphological evolution, but that remodelling of cell-cell junctions and changes in metabolism precede it.

### iPSC-derived neuroepithelial cells actively generate mechanical tension through apical constriction

While reproducible between cultures, apical areas are variable within individual cultures (Figure 1e): a hallmark of asynchronous apical constriction *in vivo*^63^. Through live imaging, we observe that iPSC-derived neuroepithelial cells spontaneously and dynamically alter their apical area (Figure 3a). Pharmacological inhibition of the myosin-activating kinase ROCK increases neuroepithelial apical area (Figure 3b-c), as previously observed in mouse embryos with equivalent ROCK inhibition *in vivo*^11^.

**Figure 3:**
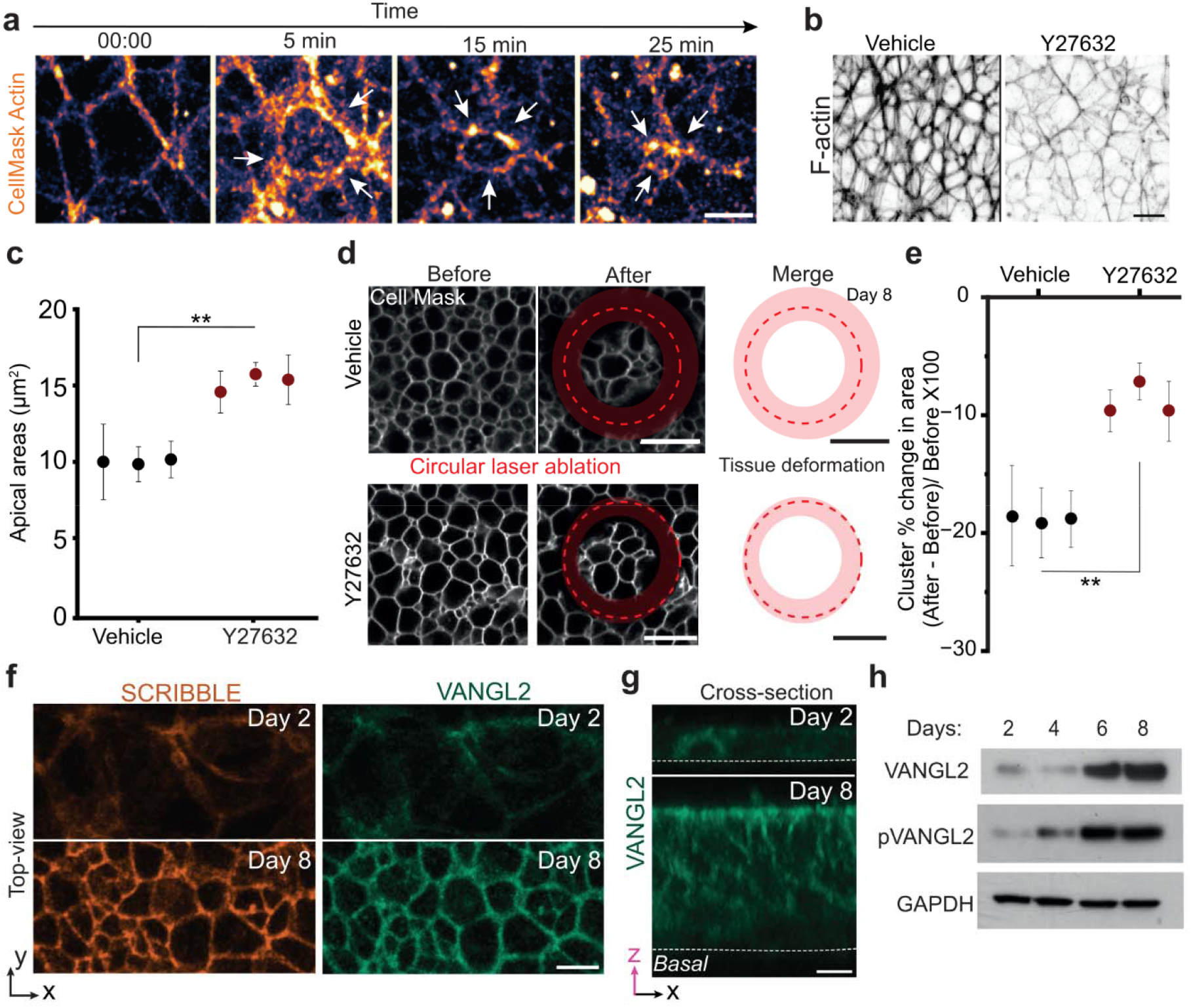
iPSC-derived neuroepithelial cells undergo ROCK-dependent apical constrictions and up-regulate expression of the PCP co-receptor VANGL2. a, Live imaging of day 8 neuroepithelial cultures stained with CellMask Actin dye. Arrows indicate the apical surface of a constricting cell (not its neighbours do not constrict). **b,** Representative F-actin-stained images of cultures treated vehicle or 10 µM ROCK inhibitor (Y27632) for 4 hr. **c,** Dot plot representing the median apical area of day 8 SFC086 neuroepithelial cultures. **d,** Live-imaged membrane-stained day 8 neuroepithelial cultures before and after a circular laser ablation, shown by red dashed lines. Red ring indicates neuroepithelial deformation before and after the ablation, respectively. **e,** Median apical cluster constriction of neuroepithelial cells, across three independent replicates in one cell line. Each point represents the median displacement quantified in each plate after at least three laser ablations. **f,g,** Representative confocal images of SCRIB and VANGL2 immunolabelled cultures at day 2 and day 8, in top-view (**f**) and cross-section (**g**). Dashed lines indicate the basal side. **h,** Representative western blot panel of VANGL2, phosphorylated VANGL2 (pVANGL2), and GAPDH protein levels across timepoints of neuroepithelial differentiation. All analyses shown are in iPSC2. P values from Wilcoxon paired test with Bonferroni correction (**: p ≤ 0.01, *: p ≤ 0.005). Scale bars = 10 μm.

Acutely expanding apical cell areas produces corrugation of the apical domain in ROCK-inhibited cultures (Supplementary Figure 3a-b), suggestive of buckling due to expansion within fixed boundaries imposed by the culture dish.

The functional outcome of apical constriction is generation of mechanical tension which is transmitted between adjacent cells and pulls on the surrounding tissue^20,64^.

Contractile neuroepithelial apical tension can be inferred through annular laser ablation assays^11,20^, in which release of a small cluster of cells from their neighbours causes rapid retraction of the cell borders within the ablated ring (Figure 3d). In our neuroepithelial cultures, the ablated surface remains within the focal plane after ablation, indicating minimal movement along the apical-basal axis. Their tension is significantly diminished following a short period of ROCK inhibition (Figure 3e), confirming that it is actively established through apical constriction.

*In vivo*, the Rho/ROCK/myosin pathway can be activated by PCP/VANGL2 signalling^20,22^, which is well-established to promote apical constriction^20,23,65^. We observe that VANGL2 and its binding partner scribbled (SCRIB) are expressed at low levels from early stages of neuroepithelial differentiation and are up-regulated by the end of the differentiation period (Figure 2f,g). Both are co-localised on the cell’s apical cortex (Supplementary Figure 4). Serine-79/82/84 tri-phosphorylation of VANGL2 also increases significantly during differentiation, with a particularly large up-regulation between days 4 and 6 (Figure 2h): coinciding with the reduction in apical area (Figure 1e) and potentially consistent with up-regulation of the ligand WNT5B (Figure 2d). Thus, iPSC-derived neuroepithelial cells express the cellular machinery required for physiological control of apical constriction and actively establish apical tension *in vitro*.

### An NTD-associated *VANGL2* mutation impairs neuroepithelial apical constriction

To our knowledge, no genetic mutations identified in human patients have previously been functionally associated with failure of apical constriction. The *VANGL2-R353C* mutation (Figure 4a) was initially reported in a fetus which had both anencephaly and spina bifida^46^, and impairs other functions of PCP signalling in fruit flies^66^. This point mutation causes a basic to polar amnio acid switch in a C-terminal region (Supplementary Figure 5) which is highly conserved across species and mediates interactions with pathway adaptor proteins^46,66^. We therefore considered it a high-priority target for testing in a human system.

**Figure 4:**
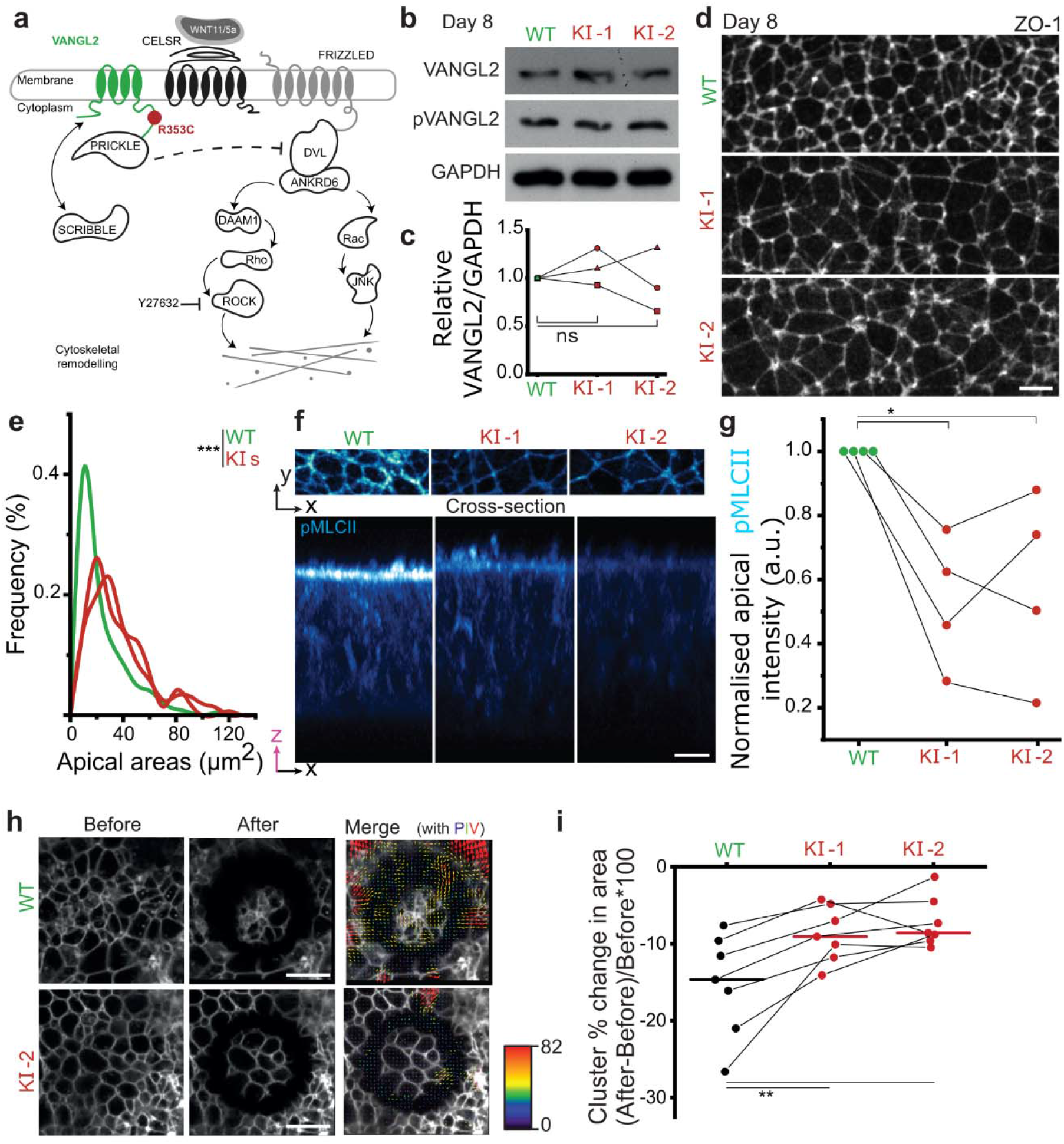
VANGL2-R353C mutation selectively impairs neuroepithelial apical constriction. a, Schematic representation of PCP pathway, starting from non-canonical Wnt ligands (e.g.Wnt5a/11) interacting with membrane co-receptors and activating cytoskeletal remodeling responses. The transmembrane protein VANGL2 interacts with CELSR, which results in downstream interaction with cytoplasmic proteins such as PRICKLE and SCRIB. The VANGL2-R353C mutation is localised at the C-terminal region of the protein, in the prickle binding domain (Pk-D). **b,** Representative western blot against VANGL2 and pVANGL2 on day 8 of neuroepithelial differentiation in the isogenic control cell line (WT) and two knock in clones with the R353C mutation (KI-1 and KI-2). **c,** Dot plots representing VANGL2 and pVANGL2 relative band intensity levels after normalisation to GAPDH, respectively, across three replicates. Intensity levels across the three lines were normalised to the WT levels of each replicate. **d,** Representative high resolution confocal images of ZO-1 immunolabelled neuroepithelial cultures on day 8, across the WT and KI cell lines. **e,** Frequency plot showing the distribution of apical area sizes of three independent day 8 neuroepithelial cultures, across the WT and both knock-in (KIs) lines. *** p < 0.001 by Kolmogorov Smirnov test, lines are the aggregate of three independent differentiations per line. **f,** Representative confocal images of neuroepithelial cultures immunolabelled for phospho-myosin light chain (pMLC)II, comparing the WT and KI-1 and KI-2 cell lines. Top view shows the distribution of protein of interest across the apical areas and cross-section illustrates localisation along the apicobasal axis. Scale bar = 10 μm. **g,** Dot plot showing normalised paired relative apical pMLCII intensity, across the WT and KI-1 and KI-2 cell lines, on day 8 of differentiation. Relative apical intensity was normalised to WT and set at 1, for each paired analysis, across four independent plates. **h,** Live-imaged membrane-stained neuroepithelial sheet before and after a circular laser ablation, with overlayed image of post-ablation timepoint and particle image velocimetry (PIV) analysis. The size and colour of arrows in PIV analysis represent the direction and magnitude of recoil. Scale = 20 μm. **i,** Median apical cluster constriction of neuroepithelial cells, across seven independent replicates. Each point represents three individual quantifications in each plate and connected lines indicate paired cultures. P values from Wilcoxon paired test with Bonferroni correction (***: p ≤ 0.001, *: p ≤ 0.005, n.s.: not significant).

We find that *VANGL2-R353C* mutation does not alter overall VANGL2 protein levels, phosphorylation (Figure 4b,c), or apical localisation (Supplementary Figure 6) in two iPSC clonal knock-in lines homozygous for this mutation compared with a congenic control line. Knock-in and congenic control lines retain their ability to differentiate into different germ layers (Supplementary Figure 7), achieve equivalent changes in neuroepithelial apicobasal thickness (Supplementary Figure 8a-c) and nuclear pseudostratification (Supplementary Figure 8d,e) during differentiation. The *VANGL2-R353C* mutation does not significantly alter cell volume, nor does it diminish extension of sub-apical protrusions which are characteristic of neuroepithelial cells^67^ (Supplementary Figure 9).

*VANGL2-R353C* knock-in lines have significantly larger apical areas (Figure 4d,e) and less apical localisation of phosphorylated myosin regulatory light chain (Figure 4f,g) than their congenic controls. Laser ablation assays confirm that apical tension is significantly lower in *VANGL2-R353C* knock-in lines than their congenic controls (Figure 4h,i). Thus, this *VANGL2* mutation associated with human NTDs selectively impairs neuroepithelial apical constriction. This is consistent with previous reports in *Xenopus* that *Vangl2* knockdown diminishes apical constriction^63,65^, but we had previously observed that mouse *Vangl2* knockout cells remain able to apically constrict when surrounded by cells which express this gene^20^. We therefore tested whether global loss of *Vangl2* impairs neuroepithelial apical constriction in a mammalian model. As expected, mouse embryos globally lacking *Vangl2* fail to close their entire spinal neural tube, developing craniorachischisis (Supplementary Figure 10a). *Vangl2*-null neuroepithelial cells show an equivalent shift towards larger apical areas in embryos (Supplementary Figure 10b-d) as seen *in vitro* (Figure 4e). Neuroepithelial apical areas are 39±8% (mean ± 95% CI) larger in *Vangl2* knockout embryos versus wild-type littermates (Supplementary Figure 10d). *VANGL2-R353C* knock-in iPSC-derived neuroepithelial apical areas were 98±32% (mean ± 95% CI across both knock-in clones with three independent experiments each) larger than their wild-type counterpart. This suggests the *VANGL2-R353C* mutation can explain all the impairment of neuroepithelial apical constriction caused by loss of *Vangl2* action.

### Establishment of patient-derived models of neuroepithelial dysfunction

As well as mechanistic and reverse genetic studies, an additional application of this highly reproducible model is patient-specific testing of key neuroepithelial behaviours (Figure 5a). Importantly, short-loop reprogramming of amniotic fluid stem cells with microfluidic delivery of modified mRNA^57^ enables rapid generation of patient-specific models in the ∼4 months between spina bifida diagnosis and birth (Figure 5a). As proof of principle, we used short-loop reprogramming to compare two iPSC lines derived from amniotic fluid cells of individuals who do not have spina bifida (here called GOC1-2 for simplicity) with two lines from individuals treated at Great Ormond Street Hospital who underwent foetal surgery for spina bifida (GOSB1-2). All four lines express comparably high levels of the iPSC marker NANOG before differentiation, and differentiate to form endoderm, ectoderm and mesoderm (Supplementary Figure 7).

**Figure 5:**
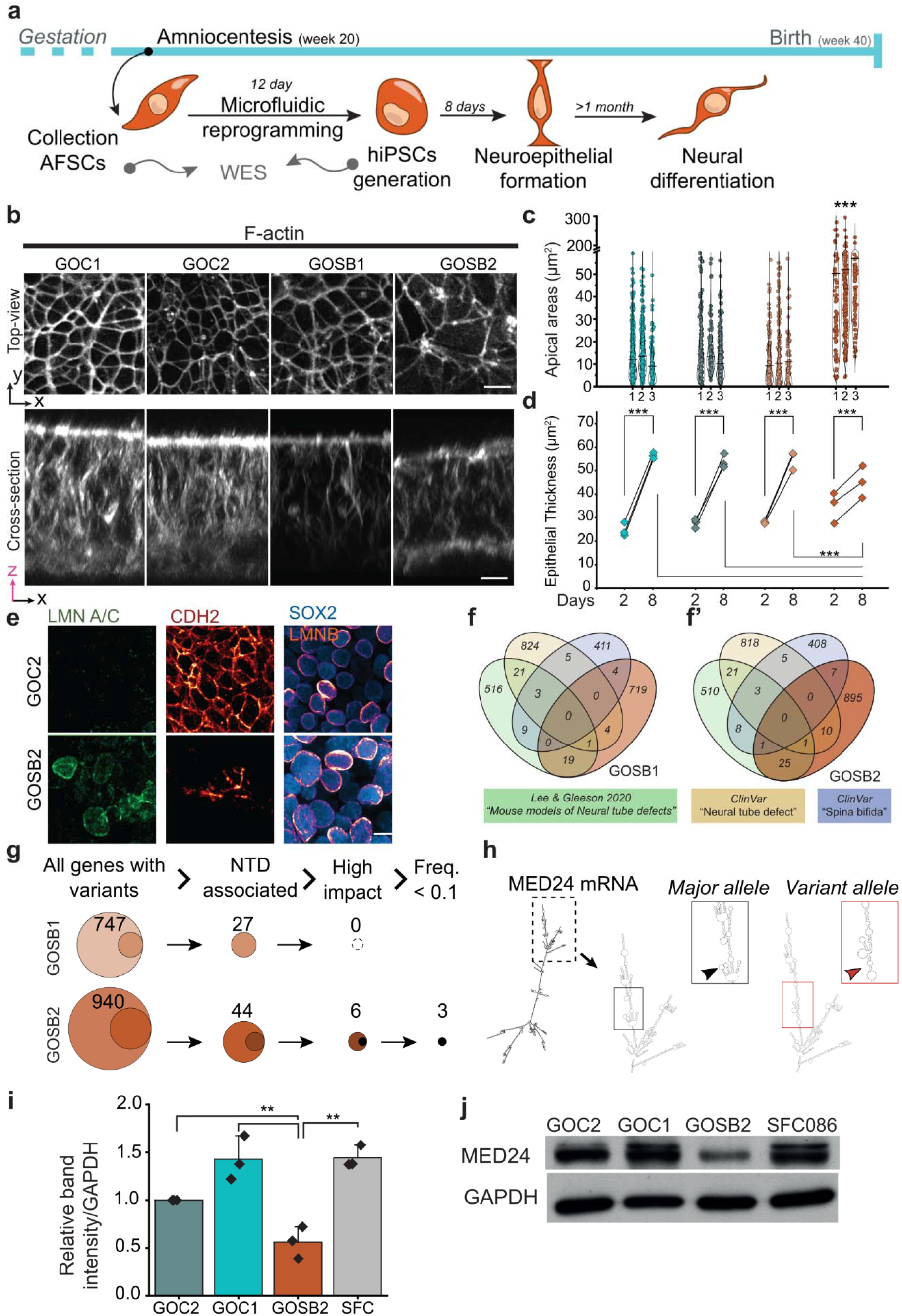
Spina bifida patient amniocyte-derived iPSC lines can identify line-specific deficiencies in neuroepithelial cell function. a, Experimental timeline of patient amniocyte-derived iPSC lines, following microfluidic reprogramming. Whole exome sequencing (WES) was performed before and after reprogramming to identify line-specific deficiencies at genomic level and pair them to neuroepithelial and/or neural function level. Initial iPSC cell expansion is not shown, but adds a few weeks to the timelines, still within the remaining period of normal gestation. **b,** Representative high-resolution top-view and cross-section images of day 8 neuroepithelial cultures visualizing F-actin. Two control cell lines (GOC1 and GOC2) were compared with two SB patient-derived hiPSCs cell lines (GOSB1 and GOSB2). **c,** Dot plot representing cell apical areas across the GOC1, GOC2, GOSB1, and GOSB2 cell lines at day 8 of neuroepithelial differentiation. Three independent plates were quantified for each line and the median apical area for each plate is shown with a black bar. **d,** Quantification of epithelial thickness on day 2 and day 8 samples, across four cell lines, GOC1, GOC2, GOSB1, and GOSB2. Points represent independent plate quantification for each line. Three paired replicates were measured for each time point and each cell line. Statistical test used for median values across quantifications was Wilcoxon paired test values with post-hoc Bonferroni. **e,** Representative confocal images of LMN A/C, CDH2, SOX2, and LMNB immunolabelling in GOC2, and GOSB2, on day 8 of neuroepithelial differentiation. **f,** Venn diagrams of GOSB1 and GOSB2 (**f’**) gene lists, after exome-sequencing, with at least one genetic variation observed in other datasets. Other datasets include previously associated mutations identified in humans under after search for the term “Neural Tube Defect” and “Spina Bifida” in ClinVar dataset, Lee and Geeson^5^. **g,** Filtering of variants in GOSB1 and GOSB2 to identify NTD-associated variants predicted to have a high impact on the transcript and which have a frequency <0.1 in gnomAD exome datasets. **h,** RNA structure of the MED24 major allele and rare variant (gnomAD reference 17-40019203-C-CACAT, allele exome frequency 0.018) predicted by RNAFold^69^. **i,** Bar plots representing normalised band intensity of MED24 across four cell lines. GOC2’s intensity levels were set as 1 in each of three replicates. **j,** Western blot panel of iPSCs from four cell lines, GOC1, GOC2, GOSB2, and SFC086-06 showing MED24 protein and housekeeping GAPDH levels. Statistical test used for values was one-way ANOVA with post-hoc Bonferroni (***: p ≤ 0.001, **: p ≤ 0.001). Scale = 20 μm.

Both control lines and the first spina bifida line -GOSB1 -achieve indistinguishable apical areas and thicknesses (Figure 5b-d), further testament to the excellent reproducibility of this modelling system when comparing non-congenic lines. In contrast, the spina bifida line GOSB2 fails to apicobasally elongate as much as other lines and retains larger apical areas (Figure 5b-d). We consider this is unlikely to represent an apical constriction defect: laser ablation assays show comparable retraction in the GOSB2 line compared with the other lines tested (Supplementary Figure 11). The epithelia formed by GOSB2 cells include mitotic cells which are not apically-located, in contrast to all other cells studied which are pseudostratified and localise their mitoses apically (Supplementary Figure 12a-d). The nuclei of GOSB2 cells are significantly larger than any of the other three lines tested at the end of neuroepithelial induction (Supplementary Figure 12e-f). Unlike all other lines tested, differentiated GOSB2 cultures also show abnormally heterogenous expression of CDH2 while partially retaining lamin A/C (Figure 5e) at the end of the standard differentiation protocol.

Attempts to adapt the protocol to ‘rescue’ this phenotype are beyond the scope of this study given the excellent reproducibility of differentiation in the seven other lines presented here.

We performed whole exome sequencing of all four amniocyte-derived lines to identify coding variants of potential clinical relevance in GOSB1 and GOSB2. Variants were compared with a list of genes previously associated with NTDs in mice and/or humans (Figure 5f, Supplementary Table 2).

GOSB1 harbours 851 variant loci within 747 genes (Supplementary Table 3; Figure 5g). Of all genes with variants, 27 are NTD-associated, such as *ZEB2* (variant 2-144389425-T-C) and *ZIC5* (13-99970996-G-T). None of the variants in NTD-associated genes are predicted to have a high transcript impact and all are heterozygous.

GOSB2, which has a more severe neuroepithelial phenotype, has 1122 variant loci in 939 unique genes of which 43 are in NTD-associated genes (Supplementary Table 4; Figure 5g). In this line, six NTD-associated genes harbour variants predicted to have a high impact on transcript sequence: *CSNK2A1*, *FAM20C*, *LFNG*, *MED24* and *PCSK5*.

Of these, *CSNK2A1* (20-503643-G-GCATATTT), *MED24* (17-40019203-C-CACAT) and *PCSK5* (9-76175222-AAATGGAATGGAATGAAATGG-A) have gnomAD variant allele frequencies <0.1. *MED24* (formerly called *TRAP100)* is of particular interest because its deletion in mice results in a thin neuroepithelium which is not pseudostratified^68^, equivalent to our observations during GOSB2 differentiation. This gene is part of the mediator complex involved in RNA polymerase II transcriptional co-activation. Our RNASeq analysis suggests *MED24* expression normally increases before neuroepithelial apicobasal elongation between days 2 and 4 of differentiation (Supplementary Figure 12g). We corroborated this increase by qPCR analysis (Supplementary Figure 12h). GOSB2 cells have compound heterozygous frameshift mutations in the same repetitive region of the final exon of this gene. We additionally confirmed that the same *MED24* variants are present in independent exome sequencing of amniocytes from which GOSB2 had been derived (not shown). Both frameshifts are in the 3’ untranslated region but are predicted to alter RNA structure, causing loss of a multi-branch loop (Figure 5h), and may therefore diminish binding of translation-regulating proteins. To test this, we compared MED24 protein levels between GOSB2 iPSCs and three control lines, finding reduced protein abundance compared to all three controls (Figure 5i,j).

As expected, given poor neuroepithelial morphogenesis of cells from the GOSB2 patient line, further differentiation results in only sparse differentiation of PAX6-positive neural progenitor cells with few neurites positive for the neuronal marker TUJ1 (Supplementary Figure 13). In contrast, GOSB1 formed prominent neuronal rosettes equivalently to the control lines (Figure 6a). However, by day 35 of neuronal differentiation, GOSB1 cultures show diminished TUJ1 positivity, without a change in the proportion of cells positive for the progenitor marker SOX2, compared with the two control lines differentiated in parallel (Figure 6b-d). Poor neuronal maturation in cells from the GOSB1 line is clearly evident by day 45 of differentiation, when control lines form dense aggregates interconnected by axons while GOSB1 cells form sparse and poorly-connected aggregates (Figure 6e,f). Thus, both the spina bifida patient-derived lines tested, in addition to *VANGL2-R353C* knock-in model, demonstrate quantifiable yet distinct abnormalities in neuroepithelial functions (Figure 6g).

**Figure 6:**
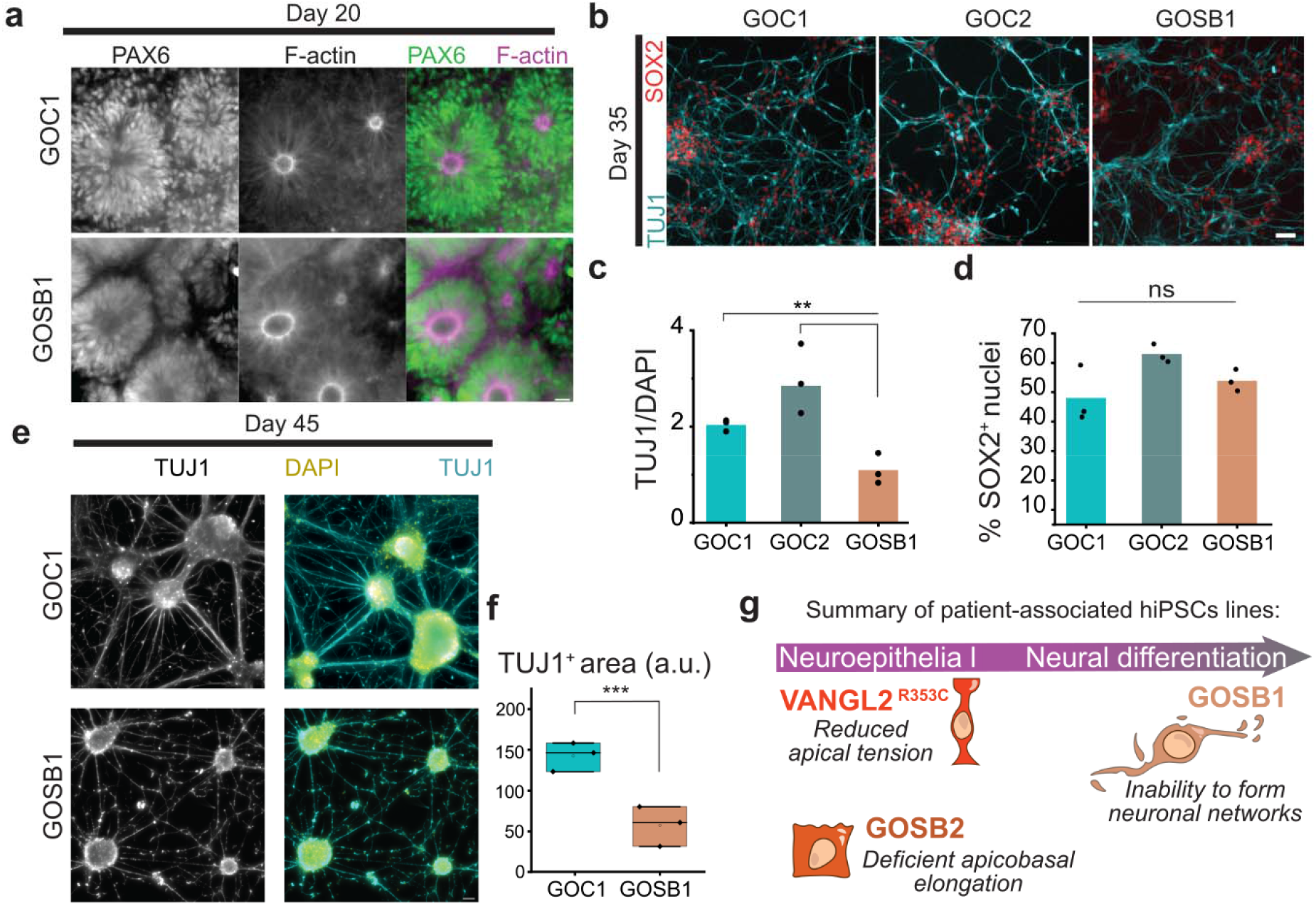
Diminished neuronal maturation suggests post-neurulation deficit of an iPSC line derived from a patient with spina bifida. a, Representative confocal images of GOC1, and GOSB1 neuronal rosette cultures (day 20) immunolabelled against PAX6 and F-actin. Scale = 100 μm. **b,** Representative immunofluorescence panel of SOX2 and TUJ1 immunofluorescence of day 30 cultures, showing GOC1, GOC2, and GOSB1 lines. Scale = 50 μm. **c, d,** Dot plot representing TUJ1/DAPI staining intensity (**c**) and the percentage of nuclei positive for SOX2 (**d**). Values were calculated in three independent replicates shown as individual dots and the average is shown with the bars. **e,** Widefield images of day 45 GOC1 and GOSB1 neuronal cultures immunolabelled for TUJ1. Scale = 200 μm. **f,** Box plot representing the TUJ1 positive area quantified across three independent cultures of GOC1 and GOSB1 cell lines, on day 45 of differentiation. P values from one-way ANOVA with post-hoc Bonferroni (***: p ≤ 0.001, **: p ≤ 0.001, n.s.: not significant). **g,** Summary of patient-associated hiPSCs lines used in this study along with the stage of failure to form a functional neuroepithelium or undergo mature neuronal formation in vitro.

## Discussion

We present a multifaceted yet tractable and reproducible model of neuroepithelial morphogenesis which is sufficiently sensitive to identify NTD-associated dysfunctions in both forward and reverse genetic studies. All three of the NTD-associated models we compare – knock-in of a patient-identified *VANGL2* mutation and two patient-derived iPSC lines – identify abnormalities which are specific to that line, and which are not present in a panel of control lines tested. The abnormalities observed relate to apical constriction, apicobasal elongation, and neuronal maturation.

Neuroepithelial apical constriction is required for closure of both the cranial and spinal neural tube^11,14,15,21^. Mouse and *Xenopus* models establish that expression of PCP signalling components enhances apical constriction^20,23,65^ but, to our knowledge, this had not previously been shown in a human system and no human genetic mutations had been linked to apical constriction as a likely pathogenic mechanism. The mechanisms by which VANGL2 promotes apical localisation and activation of myosin^20^ remain incompletely understood. They may include both PCP-specific functions of *VANGL2* equivalent to those originally identified for its orthologue in *Drosophila*, and potentially vertebrate-specific roles^70,71^. The neuroepithelial model presented here does not produce planar features commonly used as characteristic readouts of PCP signalling *in vivo*^26,40^. Nonetheless, our observation of increasing VANGL2 poly-phosphorylation suggests ligand-induced pathway activation during differentiation^35^.

This contrasts with decreasing VANGL2 phosphorylation previously reported during *in vitro* differentiation of lactogenic cells^70^.

The diminished apical constriction phenotype we document following *VANGL2-R535C* mutation occurs despite achieving wildtype-equivalent VANGL2 expression. Nonetheless, comparison with mouse embryos globally lacking *Vangl2* suggests *VANGL2-R535C* is sufficient to explain all of the dilation in apical area caused by deletion of this protein. Other mutations in the protein’s C-terminus domain which disrupt VANGL2 function sufficiently to stop neural tube closure have previously been reported in mice^20,39,40^. VANGL2’s C-terminus contains the prickle-binding domain (directly affected by the R353C mutation) and PDZ domain, which interacts with other components of the PCP pathway including SCRIB^72^. SCRIB also enhances apical constriction in mice^15,73^.

We observe that apical constriction correlates with apicobasal epithelial thickening during iPSC-to-neuroepithelial differentiation. This inter-relation is well-established in *Drosophila* ventral furrow formation^74,75^, and can be replicated following optogenetic induction of apical constriction in cultured mouse cells^76^. In mice, neuroepithelial apicobasal thickness is spatially patterned with shorter cells at the midline, under the influence of SHH signalling^14,77,78^. Apicobasal thickness of the cranial neural folds increases from ∼25 µm at E7.75 to ∼50 µm at E8.5^79^: closely paralleling the elongation between days 2 and 8 of differentiation in our protocol. The rate of thickening is non-uniform, with the greatest increase occurring during elevation of the neural folds^80^, also paralleled in our model by the rapid increase in thickness between days 4-6 as apical areas decrease. Elevation requires neuroepithelial apical constriction and these cells’ apical area decreases between E7.75 and E8.5 in mice^79^, but we and others have recently shown that this reduction is both region and sex-specific^14,81^. Specifically, apical constriction occurs in the lateral (future dorsal) neuroepithelium: this corresponds with the identity of the cells generated by the dual SMAD inhibition model we use^56^. More recently, Brooks et al^82^ showed that the rapid reduction in apical area from E8-E8.5 is associated with cadherin switching from CDH1 (E-cadherin) to CDH2 (N-cadherin). This is also directly paralleled in our human system, which shows low-level co-expression of CDH1 and CDH2 on day 4 of differentiation, immediately before apical area shrinks and apicobasal thickness increases. Also, potentially consistent with Brooks et al^82^ is our observation of down-regulation of the Sonic hedgehog effector *GLI1*, given this pathway supresses apical constriction *in vivo*.

Conservation of cells’ volume has been suggested to link apical constriction with apicobasal elongation^75^. However, we provide two lines of evidence that area/length inter-relation during neuroepithelial differentiation is not simply due to volume conservation. Firstly, cell volume decreases during neuroepithelial differentiation as thickness increases. Secondly, neither epithelial thickness nor cell volume are significantly altered in *VANGL2-R535C* neuroepithelial cells with larger apical areas.

Mechanistic separability of apical constriction and apicobasal elongation is consistent with biomechanical modelling of *Xenopus* neural tube closure showing that both are independently required for tissue bending^61^. Nonetheless, neuroepithelial apical constriction and apicobasal elongation are co-regulated in mouse models: for example, deletion of *Nuak1/2*^83^, *Cfl1*^84^, and *Pten*^79^ all produce shorter neuroepithelium with larger apical areas. Neuroepithelial cells of the GOSB2 line described here, which has partial loss of MED24, similarly produces a thinner neuroepithelium with larger apical areas.

Although apical areas were not analysed in mouse models of *Med24* deletion, these embryos have shorter and non-pseudostratified neuroepithelium^68^.

Our GOSB2 line – which retains readily detectable MED24 protein – is clearly less severe than the mouse global knockout of this gene, and the clinical features of the patient from which this line was derived are milder than the phenotype of knockout embryos^68^. Mouse embryos lacking one of *Med24*’s interaction partners in the mediator complex, *Med1*, also have thinner neuroepithelium and diminished neuronal differentiation but successfully close their neural tube^85^. Although genetic rescue experiments are beyond the scope of this study—particularly given the likelihood of gene-gene interactions—equivalence of the cellular phenotype we observed with a published mouse model, confirmation of reduced protein abundance, and preliminary evidence of gene up-regulation before apicobasal elongation which fails in GOSB2 cells, all suggest the *MED24* variants in this line are pathogenic.

As general regulators of polymerase activity, MED proteins have the potential to alter the timing or level of expression of many other genes, including those already known to influence pseudostratification or apicobasal elongation. MED depletion also causes redistribution of cohesion complexes^86^ which may impact chromatin compaction. MED family proteins are increasingly implicated in human neural tube defects: *MED12* and *MED13L* likely pathogenic variants have been reported in patients^9,87^. Their functions during neural tube closure, including through known interactions with Wnt and PCP pathways^88^, remain incompletely understood and future work will be needed to determine whether *MED24* mutation is the only genetic change relevant to neuroepithelial morphogenesis in GOSB2. MED protein contributions to neurogenesis and neuronal maturation are more well-established^85,89–91^.

Another limitation of our model, shared by many *in vitro* systems, is the poor ability to replicate maternal environmental influences. For example, the validated cell culture media used for this model contain 3.4 mg/L folic acid instead of folate. The down-regulation of one-carbon metabolism-associated transcripts we observe before apical constriction was unexpected because exogenous folic acid can trigger apical constriction^92,93^. Folic acid in the medium may mask NTD predispositions predisposed by maternal folate deficiency or genetically impaired placental transport, while accentuating cellular deficiencies which diminish folic acid conversion into a one-carbon donor^94^. We cannot exclude *in vitro* accentuation of the cellular phenotype of the GOSB2 line, as it is difficult to imagine that such severe failure of neuroepithelial morphogenesis would produce a sufficiently normal spinal cord to benefit from fetal surgery performed on the patient these cells were derived from. Alternatively, compensation may be possible *in vivo*, potentially through interactions between tissue types which promote robustness of morphogenesis.

Some of these limitations, potentially including inclusion of environmental risk factors, can be addressed by using alternative iPSC-derived models^93,95,96^. For example, if patients have suspected causative mutations in genes specific to the surface (non-neural) ectoderm, such as *GRHL2/3*, 3D models described by Karzbrun et al^49^ or Huang et al^97^ may be informative. Characterisation of surface ectoderm behaviours in those models is currently lacking. These models are particularly useful for high-throughput screens of induced mutations^97^, but their reproducibility between cell lines, necessary to compare patient samples to non-congenic controls, remains to be validated. Spinal cell identities can be generated in human spinal cord organoids, although these have highly variable morphologies^98,99^. As such, each iPSC model presents limitations and opportunities, to which this study contributes a reductionist and highly reproducible system in which to quantitatively compare multiple neuroepithelial functions.

A key benefit of our system is the natural progression from cell functions relevant to neurulation, to neurogenesis. Others previously reported that iPSC lines derived from individuals who have spina bifida are less able to differentiate into neurons than those from healthy controls^100^. This is consistent with our findings in the GOSB1 patient-derived line. Although gene rescue experiments are beyond the scope of this study, potentially deleterious variants identified in the GOSB1 patient line, including a *ZIC5* variant, may impair neuronal development. The reduction in TUJ1 positive cells observed in this line compared with non-congenic controls is at a point in differentiation which is highly comparable between individuals^101^. Individuals with spina bifida commonly have thinning of the neocortex and other supratentorial brain abnormalities^102^. What is less clear is whether diminished neurogenesis, which normally occurs after closure of the neural tube, may relate to causation of NTDs. The converse – that open spina bifida impairs neural development – is directly supported by experimental evidence^103^. Nonetheless, impaired neuronal differentiation of patient-derived iPSC lines raises the potential for shared genetic/epigenetic causation of the spinal lesion and cortical brain abnormalities, potentially compounded by abnormalities in cerebrospinal fluid drainage and amniotic fluid exposure.

In summary, our findings in a human neuroepithelial model corroborate animal studies identifying diminished apical constriction as a cause of NTDs, implicate VANGL2 acting upstream of myosin phosphorylation in this behaviour, identify diminished neuroepithelial thickness in a patient-specific disease model and corroborate the importance of MED family proteins known to enhance this process as an emerging genetic contributor to human NTDs. The multi-genic nature of neural tube defect susceptibility, compounded by uncontrolled environmental risk factors (including maternal age and parity^104^), mean that patient-derived iPSC models are unlikely to provide mechanistic insight. They do provide personalised disease models which we anticipate will enable functional validation of genetic diagnoses for patients and their parents’ recurrence risk in future pregnancies, and may eventually stratify patients’ postnatal care. We also envision this model will enable quality control of patient-derived cells intended for future autologous cell replacement therapies, as is being developed in post-natal spinal cord injury^105^. Thus, the highly reproducible modelling platform we evaluate – which is robust to differences in iPSC reprogramming method, sex and ethnicity – represents a valuable tool for future mechanistic insights and personalised disease modelling applications.

## Materials and Methods

Reagents and key resources are tabulated in Appendix 1.

### Human iPSC culture maintenance

All iPSCs lines were grown in E8 medium (ThermoFisher, A1517001), using 0.5% Matrigel Growth Factor Reduced (Corning, 354277) coated plates, according to a previously published protocol^106^. Cells were routinely grown in 6 well plates (SARSTEDT, 833920300), unless otherwise stated. E8 medium was fully replaced daily. At ∼80% confluency cells were washed with PBS without CaCl_2_/MgCl_2_ (ThermoFisher,14190094) and dissociated with 0.5 mM EDTA (ThermoFisher, 15575020) for 5 min. Cells were resuspended in E8 medium and re-plated at a 1:10 split ratio. All cell lines were tested monthly for mycoplasma contamination, using the MycoAlert^®^ kit (Lonza, LT07-318). A summary with the characteristics and reprogramming technologies of the hiPSCs lines used on this study can be found in Table 1.

**Table 1:**
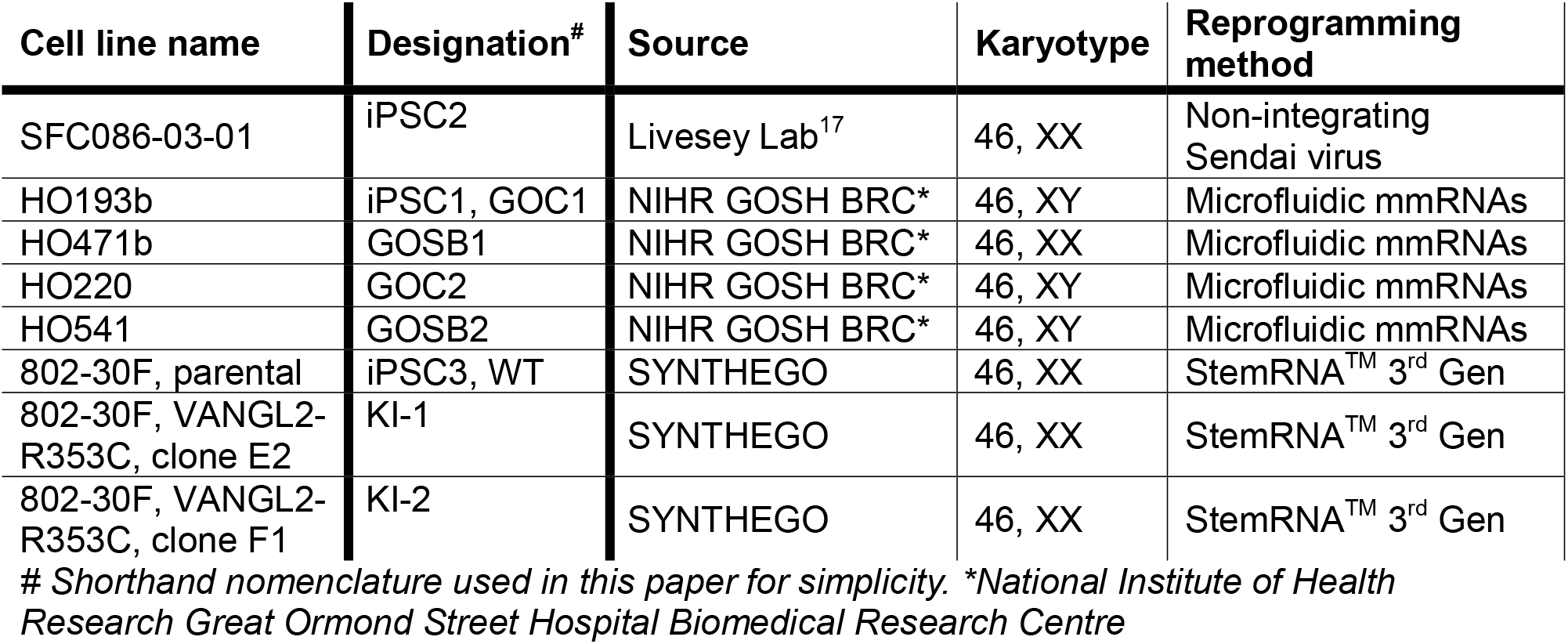
hiPSCs line information.

Unless otherwise stated, for each iPSC line three independently differentiated plates were generated and analysed, with each plate representing a separate differentiation experiment performed on different days. For patient-specific iPSC lines, one independent iPSC line was obtained per patient following microfluidic mmRNA reprogramming^57,107^. Amniocyte reprogramming was performed as a service by the NIHR Great Ormond Street Hospital Biomedical Research Centre under ethical approval IRAS number 133888, REC number 14/LO/0863 (Title “Amniotic fluid, placental and fetal stem cells”).

### VANGL2 knock-in cell line generation

The selection of the mutation was made according to a previously identified gene alteration in a patient described as having anencephaly with occipital and cervical spina bifida^46^. The specific point mutation was found in heterozygosity and was located on chromosome 1, at nucleotide 160,421,174 in the genomic sequence (Supplementary Figure 5). Although other *Vangl2* mutations have been found to exert dominant negative effects^108^, we generated homozygous mutant lines to facilitate interpretation of results obtain. In collaboration with SYNTHEGO, an established hiPSC line 802-30F was genetically engineered to introduce a homozygous point mutation by replacing a cytosine (C) with a thymine (T) (CGC>TGC), at a specific genetic locus. After high knock-in efficiency was confirmed, specific sgRNA with Cas9 were electroporated to the cells. VANGL2 knock-in lines were generated using CRSIPR-Cas9 homology directed repair editing by Synthego (SO-9291367-1). The guide sequence was AUGAGCGAAGGGUGCGCAAG and the donor sequence was *CAATGAGTACTACTATGAGGAGGCTGAGCATGAGCGA AGGGTGTGCAAGAGGAGGGCCAGGTGGGTCCCTGGGGGAGAAGAGGAGAG*.

Sequence modification was confirmed by Sanger sequencing before delivery of the modified clones, and Sanger sequencing was repeated after expansion of the lines (Supplementary Figure 5) as well as SNP arrays (Illumina iScan, not shown) confirming genomic stability. For the generation of the isogenic control line (WT), the cells were electroporated with SpCas9 only and confirmed to be unedited at target locus. Approximately two weeks later the cells colonies have expanded and genotyping of the pool of clones was undertaken. Individual clones were then isolated and clonally expanded to confirm the presence of the knock-in genetic alteration. Those clones positive for the genetic alteration were further Sanger sequenced to confirm stable *VANGL2* editing.

### Adherent neural induction protocol

Direct differentiation of hiPSCs to neuroepithelial sheets was as we previously reported^17^. Briefly, hPSCs were lifted with EDTA and plated at 100% confluency, ∼300,000 cells/cm^2^, in E8 medium. It is critical to ensure that 100 % confluency is reached at seeding (considered ‘day 1’). After 24 hours, cells were washed twice with PBS without CaCl_2_/MgCl_2_ and the medium was replaced with 2 ml of neural maintenance media (NMM, see Table 2), supplemented with 1 μM Dorsomorphin (Tocris, 3093) and 10 μM SB431542 (Tocris, 1614). NMM with these supplements is referred as neural induction medium (NIM) and was refreshed daily until day 8. At that timepoint the cultures were terminated or passaged for neuronal differentiation.

**Table 2:** Composition of neural maintenance medium, at 50:50 mixture.

| Product | Volume (ml) | Company | Cat. Number | Storage (°C) |
| --- | --- | --- | --- | --- |
| **Solution A (500ml):** |  |  |  |  |
| 2-Mercaptoethanol | 1 | Gibco | 21985-023 | 4 |
| DMEM/F12+Glutamax | 481.25 | Gibco | A15169-01 | 4 |
| Insulin | 0.25 | Sigma | 19278 | 4 |
| N2 supplement (100x) | 5 | Gibco | 17502-048 | -20 |
| Non-Essential amino Acids | 5 | Gibco | 11140-050 | 4 |
| Penicillin/Streptomycin | 2.5 | Gibco | 15140-122 | -20 |
| Sodium Pyruvate | 5 | Sigma | S8636 | 4 |
| **Solution B (500ml):** |  |  |  |  |
| B27-supplement | 10 | Gibco | 17504-044 | -20 |
| L-Glutamine | 5 | Gibco | 25030-024 | -20 |
| Neurobasal | 482.5 | Gibco | 12348-017 | 4 |
| Penicillin/Streptomycin | 2.5 | Gibco | 15140-122 | -20 |

Neuronal differentiation was performed as previously described^56^. The neuroepithelial sheets were passaged at day 8, using Dispase (ThermoFisher, 17105041) for 15-20 minutes at 37°C. After washing with PBS, neuroepithelial clumps were plated on 1% Matrigel-coated 6-wells in NIM, ensuring sufficiently sparse seeding to give clumps space to form rosettes. After 24 hours of passaging, the medium was changed to NMM supplemented daily with 20 ng/ml of FGF2 (PeproTech, AF-100-18B) for the next four days. After day 12, neural rosettes were passaged to 1% Matrigel coated plates, using Dispase. NMM was then changed every 48 hours. This process was repeated at least three-times, when confluency reaches 80 %, until day 25 when the neural progenitor cells mature. At that stage, once the confluency reached 80 %, cells were passaged with Accutase (Sigma, A6964) to dissociate the rosettes into single cells. At that stage (∼day 30) cells can be cryopreserved or plated for the last time at 25,000-75,000 cells/cm^2^ on Laminin-521 (Gibco, A29249) coated plates.

### Bulk RNA-seq and Cross-Correlation Analysis

Bulk RNA sequencing was performed to characterise in detail the transcriptional changes during neuroepithelial induction of iPSC3 cells. 100 ng of total RNA was extracted using the RNeasy Mini Kit (Cat. No. 74104, Qiagen) according to the manufacturer’s instructions from five culture wells per timepoint and pooled into a single sample per timepoint. RNA quality and concentration were assessed prior to sequencing. Five independent biological replicates were generated for each time point across days 1–8 of neuroepithelial induction.

RNA samples were submitted to GeneWiz (Azenta Life Sciences) for library preparation, quality control and high-throughput sequencing. RNA integrity was assessed by agarose gel electrophoresis and RNA concentration was determined using a Qubit Fluorometer (Thermo Fisher Scientific). Library preparation, sequencing and primary data processing were performed by GeneWiz using their standard bulk RNA-sequencing pipeline. Raw sequencing reads were processed through GeneWiz’s bioinformatics workflow, and the resulting gene expression matrices were used for downstream analyses.

Cross-correlation analysis was performed by first identifying all the genes which underwent at least a two-fold change in expression between any two timepoints in their differentiation trajectory (max/min expression ≥2). Next, the linear correlation coefficient was calculated between the expression of those genes versus apical area quantified in parallel cultures. A slope **p** value was calculated for each gene based on the **t**-statistic for each correlation **r** value. The resulting **p** values were then corrected for multiple comparisons using the False Discovery Rate (FDR) method. Cross-correlation was then performed by comparing correlation **r** values between gene expression versus apical area in synchrony (expression on days 2-8 versus area on days 2-8), when expression is offset by +1 day (expression on days 3-8 versus area on days 2-7), or when expression is offset by -1 day (expression on days 2-7 versus area on days 3-8). Genes where then separated according to the correlation which produced the largest **r** value and processed by gene ontology (GO) analysis using ShinyGO 0.80^109,110^.

### Generation of *Vangl2* knockout mouse embryos

Studies were performed under the regulation of the UK Animals (Scientific Procedures) Act 1986 and the National Centre for the 3Rs’ Responsibility in the Use of Animals for Medical Research (2019). Breeding of transgenic mice was performed under Home Office license with local Animal Welfare Ethical Review Board approval. Mice were mated overnight, and the next morning was considered to be E0.5 if a plug was found. *Vangl2* knockout mouse embryos were as previously described^111^.

### Immunofluorescent staining and confocal microscopy

Adherent cultures were stained for a maximum of 20 minutes in cold 4% paraformaldehyde (PFA) and immunofluorescent staining as previously described^17^. Primary antibodies used were rabbit anti-ROCK (Abcam antibody ab45171), rabbit anti-Ser19 pMLC (Cell Signalling Technology antibody #3671), rabbit anti-SOX2 (Merck antibody ab5603), mouse anti-CDH2 (Cell Signaling Technology antibody #14215), mouse anti-Lamin A/C (Santa Cruz antibody sc7292), goat anti-Lamin B1 (Santa Cruz antibody sc6216), rabbit anti-ZO1 (Thermo Fisher Scientific antibody #40–2200), goat anti-scribble (Santa Cruz antibody sc-11049), rat anti-VANGL2 (Merck antibody, MABN750), rabbit anti-Pax6 (Biolegend antibody 901301), and mouse anti-TUJ1 (BioLegend MMS-435P), and detected with Alexa Fluor™ conjugated secondary antibodies (Thermo Fisher Scientific). All primary and secondary antibodies were used in 1:200 and 1:500 dilution, respectively. Nuclei were labelled with DAPI (Severn Biotech Ltd 17057) and F-actin was labelled with Alexa Fluor™-647 conjugated phalloidin (Thermo Fisher Scientific). Confocal salt and pepper noise was removed using a ‘remove outliers’ filter and linear adjustments to brightness and contrast were applied to all parts of each figure panel equally.

Confocal images were captured on a Zeiss Examiner LSM880 confocal, using a 20x/NA1.0 Plan Apochromat dipping objective with AiryScan Fast. Widefield images of the neural rosettes and neuronal cultures were captured on a Zeiss Observer inverted using 10x/NA0.45 objective or a Nikon Ti2 Eclipse widefield microscope using 10x/NA0.30 or 20x/NA0.45 Plan Apochromat λD objectives.

### Laser ablations and live imaging

Annual laser ablations were performed as previously reported^11,20,112^ on day 8 neuroepithelial sheets. Briefly, cells were homogenously stained with CellMask membrane dye (Invitrogen, C10046) and a single apical z-slice was selected for the target region. The ablation circle was randomly placed in the stained neuroepithelial sheet and three ablations in different fields of view were performed per dish then averaged (no predictable change in tension was observed between repeated ablations). The percentage reduction in cell area within the ablated annulus immediately after ablation was quantified. The imaging order for each cell line was alternated between each independent replicate to ensure no bias in the analysis and recoils were analysed blinded to genotype/treatment.

For live-imaging sessions, day 8 neuroepithelial sheets were stained for 5 minutes with CellMask Deep Red Actin Tracking stain (ThermoFisher, A57245) in NIM. Imaging was performed with the Zeiss LSM880 confocal microscope using 20x/NA1.0 Plan Apochromat dipping objective. The microscope chamber was pre-warmed at 37 °C and CO_2_ was set at 5 %. The water-immersive objective was cleaned with 70 % ethanol solution and let to air dry to remove any potential contaminants. A plexiglass chamber was assembled around the culture plate to maintain stable culture condition. Damp cotton wool was put in the plexiglass set up to create a humidified chamber and avoid media evaporation. ZEN Black v2.3 was used to operate the microscope and capture images. Apical region images were acquired every 5 minutes.

### Particle intensity velocimetry (PIV) analysis

Particle Image Velocimetry (PIV) examined the flow of apical areas of day 8 neuroepithelial sheets after laser ablations. Briefly, we identify changes in pixel flow before and after the ablation. For the PIV analysis used in this report a pre-installed plug in FIJI was used^113^. The analysis considers the displacement of pixels within a specific region, termed ‘interrogation window’. This area in the present study was specified between two frames, before and immediately after the circular laser ablation. For the analysis a ‘Pure Denoise’ filter (Cycle-spins: 4, Multiframe: 2) was used to ensure that non-specific random noise is not interpreted as movement.

### Quantitative cell morphology analysis

Apical area analyses were performed in F-actin stained neuroepithelial sheets, using the polygon selection tool in FIJI and outlining the apical cap of individual cells and saving them as ROIs. At least 100 contiguous cells were analysed from each image.

Epithelial apicobasal thickness, confocal images were optically resliced and their cross-section outline manually annotated. Thickness was calculated using the ‘bounding rectangle’ (width) tool in ImageJ/Fiji. This was performed in three images across each dish and the average height was calculated for each culture.

For fluorescent signal polarisation analysis, the plot profile command in ImageJ/FIJI was applied to an optically-resliced section of equal thickness within each dataset.

### TUJ1 and SOX2 neuronal quantification

For the quantification of TUJ1+ axons of neural progenitor, the total area of axons was quantified relative to the area imaged. The TUJ1 signal was binarized using the ‘Threshold’ command, and the total area of the axons was summed. Finally, the TUJ1+ area was normalised to the total number of cells per field of view.

For the quantification of SOX2 nuclear marker in neuronal progenitor cultures, a multi-step pipeline was used. Initially, all nuclei were identified in the DAPI-stained channel and using the command ‘Find Maxima’ in FIJI. A binarized image was generated, with individual points within each nucleus. Each point was enlarged and used to generate a ROI within each nucleus The individual points were used as a reference to quantify the SOX2 intensity within each cell. A threshold was set manually after identifying the nucleus with the lowest positive signal. Finally, the percentage for positive nuclei was calculated after subtracting the total cell numbers with the positive proportion and dividing by the overall cell number.

### Western blotting and quantification

Western blotting was performed as previously described^114^. Neuroepithelial cells were lysed on ice for 30 min in 50 mM Tris Base pH 7.6, 150 mM NaCL, 1% Triton X-100, 0.02% Sodium Azide, 1x protease inhibitor cocktail (Merck Life Science UK Ltd), 1 mM sodium orthovanadate, 25 mM sodium fluoride. The protein concentration was determined using Pierce BCA Protein Assay kits (Thermo Fisher). Proteins were resolved by SDS-PAGE using 10-12% polyacrylamide gels and transferred to PVDF membranes. Membranes were blocked in TBS (15.4 mM Trizma-HCL, 4.62 mM Tris-base, 150 mM NaCl, pH 7.6) containing 10% w/v milk powder (Merck Life Science UK Ltd). Membranes were incubated with primary antibody overnight and then with horseradish peroxidase (HRP)-conjugated secondary antibodies (Agilent Technologies, Stockport, UK) for 1 hour. Primary antibodies were used rabbit anti-pVangl2 (Merck antibody SAB5701946), mouse anti-NANOG (Abcam antibody ab173368), rabbit anti-OCT4 (Abcam antibody ab19857), mouse anti-CDH1 (R&D FAB18381A), mouse anti-GAPDH (Merck antibody MAB374), rabbit anti-MED24 (BETHYL antibody A301-472A). HRP was detected using ECL Prime (Cytiva, Amersham, UK). To strip and re-probe, membranes were incubated in 0.2 M NaOH for 20 min at 37 °C and then 20 min at room temperature before re-blocking and re-use.

For the quantification of western blot films, the band area method was applied using ImageJ/FIJI. A box was drawn around the bands and after ensuring all bands fit within it, and the intensity profile was plotted. The total area between the line graph and x axis, after removing background, was quantified for each sample. The protein level was normalised to the housekeeping protein, LMNB or GAPDH. The normalised protein levels were compared between samples, and a control value was normalised to 1. Protein quantifications were performed in paired replicates or in three independent cell lines.

### Whole exome sequencing

Approximately 500 ng of DNA from each cell line, namely GOC1, GOC2, GOSB1, and GOSB2, were purified. The subsequent steps of fragmentation, library preparation, Illumina sequencing were performed using established pipelines by Azenta, UK. Upon receipt of the samples, a quality control (QC) screening was executed, and all samples successfully pass the QC threshold including concentration and maintaining high DNA integrity number (DIN, >7.8). FASTQ data QC, genomic alignment, variant colling and annotation was also performed by Azenta, UK. For all samples, >97.5% of target regions achieved a read depth ≥20X. Variants were annotated using the Sequence Ontology consensus terms. The following were considered to have a high impact: transcript_ablation (SO:0001893), splice_acceptor_variant (SO:0001574), splice_donor_variant (SO:0001575), stop_gained (SO:0001587), frameshift_variant (SO:0001589), stop_lost (SO:0001578), start_lost (SO:0002012), and transcript_amplification (SO:0001889).

### Reverse-transcription quantitative PCR (RT-qPCR)

Total RNA was isolated from neuroepithelial samples using the RNeasy Plus Mini Kit (Qiagen, 74134) following the manufacturer’s protocol. Reverse transcription was performed with 200 ng of RNA using the ProtoScript® II First Strand cDNA Synthesis Kit (New England Biolabs, E6560L) as per manufacturer’s instructions. Quantitative PCR was carried out using the Luna® Universal qPCR Master Mix (New England Biolabs, M3003L) in 10 µL reactions containing 2 ng of cDNA and 0.5 µM of forward and reverse primer. Fluorescence was detected using the Bio-Rad CFX96 C1000 Touch Real-Time PCR Detection System with all samples measured in triplicate and values normalised to *GAPDH*, the housekeeping control. Relative expression of *MED24* compared a control timepoint was calculated using the 2-ΔΔCt method. Primer sequences: *GAPDH* (For – TTGGCTACAGCAACAGGGTG, Rev – GGGGAGATTCAGTGTGGTGG), *MED24* (For – TCTCATGGAGACAAGGACTTCA, Rev – CGCTTAGCCATAAGGTTGTTGA).

### NTD associated gene sorting workflow

Lists of genes with annotated variants in GOSB1 and GOSB2 were compared with GOC1 and GOC2. Only variants not shared with GOC1/2 were considered in order to eliminate batch effects and very common variants. Each of resulting gene lists was then cross-referenced with the most current compilation of mouse models associated with NTDs, as documented in Lee and Gleeson^5^. Two additional gene lists were generated based on searches in ClinVar database for entries linked to “Neural Tube Defect” and “Spina Bifida” (Supplementary Table 2). Collectively, these identified a subset of variant genes associated with NTDs. These were subsequently filtered by sequency ontology severity score, and the allele frequencies of high impact variants was determined using gnomAD v4.1.0^115^.

### Statistical analysis and reproducibility

Comparisons between two groups were by Student’s t-test accounting for homogeneity of variance in SPSS (IBM Statistics 25). Comparison of multiple groups was first tested for normality and homogeneity of variance in Origin 2020 (Origin Labs). According to the result if samples were normally distributed an ANOVA test was performed, while for non-parametric datasets a Kruskal-Wallis ANOVA was used in Origin 2020 (Origin Labs).

Post-hoc pairwise analysis was performed to determine the statistical significance. P values <0.05 were considered statistically significant. All graphs were generated in Origin 2020. Replicates were defined as technical replicates (same iPSC line cultured at the same time) or biological replicates (different iPSC lines or same line cultured independently). In the case of VANGL2 knock-in lines, two clones were compared with their congenic control line. To avoid bias, blinding and randomisation was performed were possible by asking a lab member to rename copied files, without disclosing cell line information prior to analysis. However, changes of neuroepithelial thickness across time points days were too dramatic to not know which day was analysed and morphological differences of GOSB3 also precluded blinding.

## Data availability

All data presented in this manuscript is included in the main figures, supplementary files, source data Excel file and uncut western blot PowerPoint file.

## Acknowledgements

The authors would like to thank Prof F Livesey and Dr F Paonessa for their assistance in establishing the dual-SMAD inhibition protocol in our lab and provision of SFC086-03-01 as a reference line. We thank Jessica Reeves for technical assistance with western blots, Louis Hamilton for help with apical area segmentation, Dr Teresa Pereira da Silva for scientific discussions and cell culture assistance, Dr Timo N. Kohler and Clara Munger for their critical reading of the manuscript, and Dr Dale Moulding for assistance with imaging. IA was funded by the NIHR GOSH Biomedical Research Centre. Views expressed in this manuscript are solely those of the authors and not necessarily those of the NHS, the NIHR or the Department of Health. GLG acknowledges funding from the Royal Society (RG\R2\232082), Rosetrees (2023\100266), NC3Rs (NC/Y50063X/1) and Wellcome Trust (211112/Z/18/Z).

## Author contributions

IA, EMT, YG, AK and GLG performed and analysed experiments. GLG, PDC and NE conceived the study and designed experiments with IA. GLG, PDC and IA secured funding. GLG, PDC, NE and EM supervised staff. GLG and IA wrote the manuscript with input from all co-authors.

