## Supplementary Figure for "Patient-specific iPSC models of neural tube defects identify deficiencies in neuroepithelial cell shape regulation and differentiation"

### Supplementary Figures and Legends

Supplementary Figure 1: Evolution of cell shapes during neuroepithelial differentiation..2

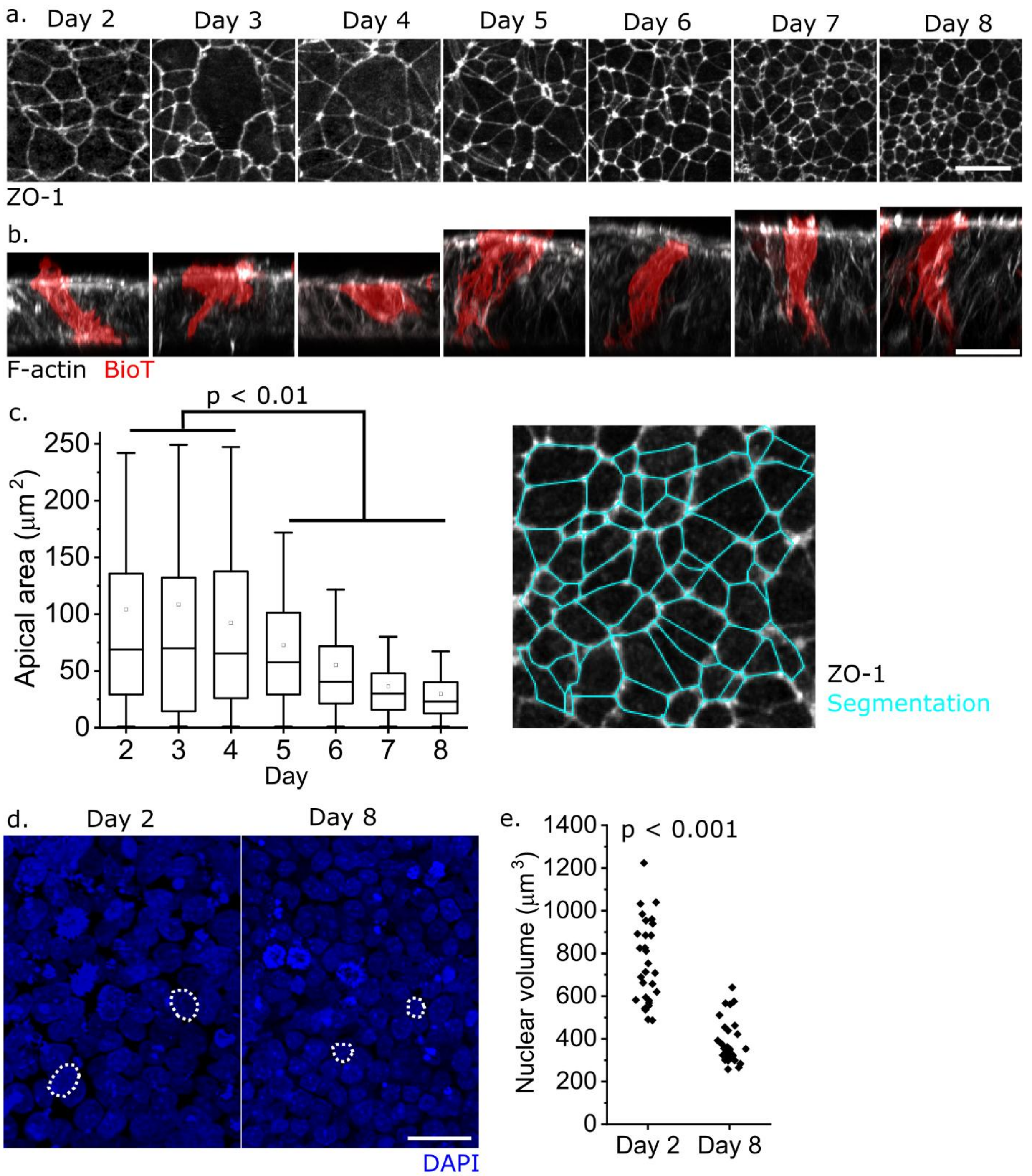

**Supplementary Figure 1: Evolution of cell shapes during neuroepithelial differentiation**. ***a,*** *ZO-1 immunofluorescence showing the apical surface of differentiation iPSC3 cells between day 2 (24 hours after initiation of neural induction medium treatment) and day 8*. ***b,*** *Lateral views of iPSC3 cells over the course of differentiation showing the rapid increase in apicobasal length after day 4 and complexity of cell shapes. Scale bars = 25 µm, signal intensity at each timepoint adjusted individually.* ***c,*** *Quantification of apical area (mean ± 95%CI) based on ZO-1 segmentation of iPSC3 cells fixed between days 2-8 of neuroepithelial induction (one replicate,*

*boxes show median and inter-quartile range with whiskers showing outlier range). P values by ANOVA with Bonferroni post-hoc indicate days 2-4 are not significantly different from each other but all are significantly larger than days 5-8.* ***d,*** *Visualisation of nuclei on day 2 and 8 of neuroepithelial differentiation. Dashed white lines indicate nuclear outlines. Representative of multiple iPSC lines including iPSC1-3. Scale = 25 µm. e, Quantification of nuclear volume measured by manually drawing around each nucleus in AiryScan Z-stacks spread throughout the apicobasal thickness of the epithelium. Points represent individual nuclei (30 nuclei per day, 10 nuclei from three differentiation experiments).*

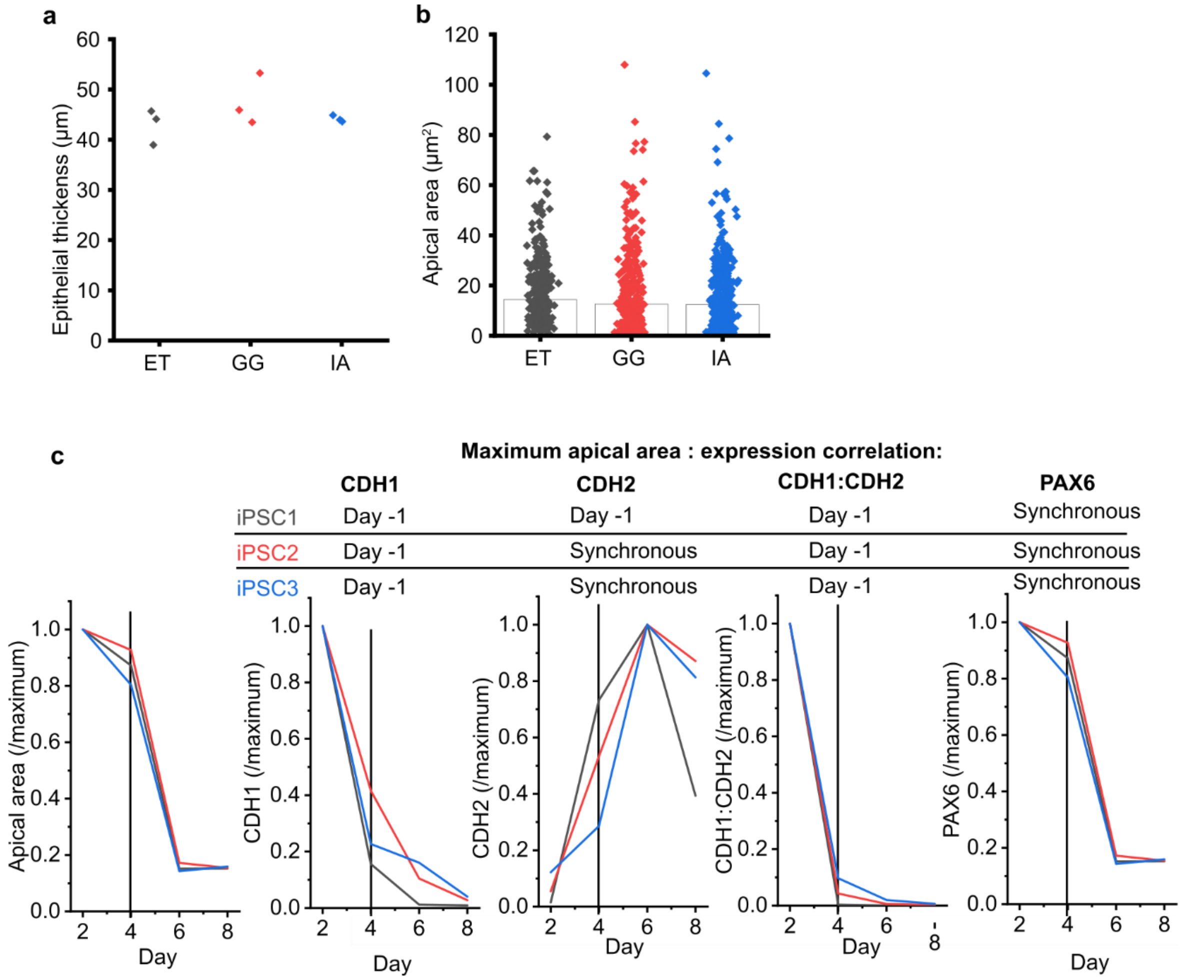

**Supplementary Figure 2: Comparison of neuroepithelial morphology in cultures differentiated by different investigators and correlation with expression of selected markers. a-b**, *Neuroepithelial (day 8) thickness (**a**) and apical area (**b**) in iPSC2 cells differentiated by ET, GG and IA. All were analysed by GG. Points represent different cultures (**a**) and individual cells (**b**, 300 cells from 3 cultures per investigator, with bars showing medians). No groups are significantly different from others by T-test.* ***c,*** *Plots of changes in apical area and quantification of selected markers (see Figure 1) used for cross-correlation analysis. The vertical lines indicate day 4: apical area changes minimally before this,*

*whereas cadherin expression changes earlier in most cases. Cross-correlation analysis is performed by comparing the correlation between protein abundance and median apical area (divided by the maximum value, such that this is normalised to 1). Cross-correlation identifies the maximum correlation coefficient (r value) when the two parameters are temporally shifted relative to each other. 'Day -1' means that correlating protein levels on day 4-8 versus apical area on days 2-6 produces a higher r value than when protein and area are correlated across days 2-8 (synchronous).*

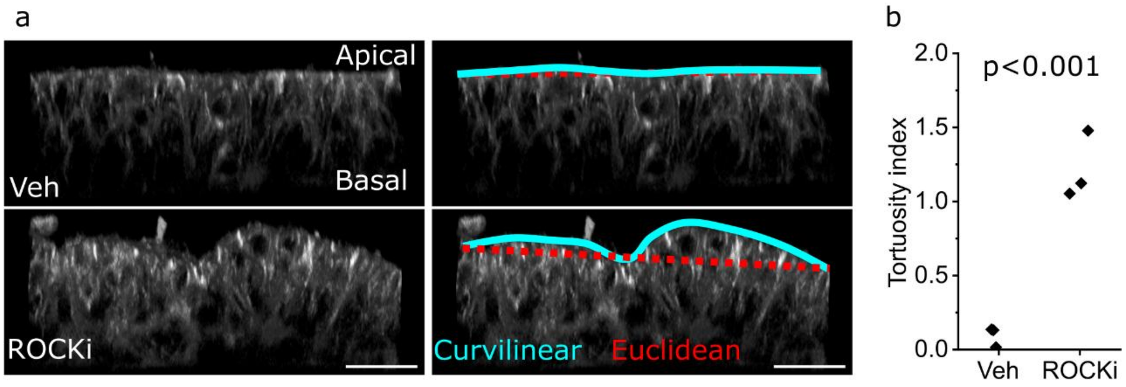

**Supplementary Figure 3: Neuroepithelial buckling following apical expansion.** ***a,*** *Lateral views showing 3D reconstructions of the neuroepithelium (day 8) in vehicle-treated and ROCK-inhibited cultures. The cyan lines indicate to total curvilinear length of the apical surface across the field of view with a length indicated by the red Euclidean distance. Scale bars = 50 µm.* ***b,*** *Quantification of apical surface tortuosity defined as (Curvilinear – Euclidean)/Euclidean * 100. P value from T-test, points represent independent cultures.*

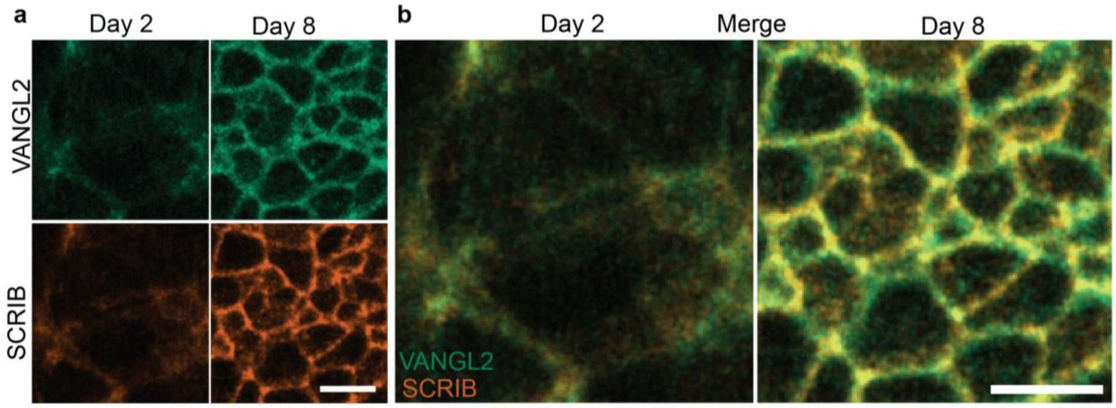

**Supplementary Figure 4: SCRIB and VANGL2 co-immunolocalization in control hiPSCs.** ***a.*** *Representative high-resolution max projection images of day 2 and day 8 neuroepithelial cultures stained against SCRIB, and VANGL2, in a control 802-30F line.* ***b.*** *Overlay of images showing co-immunolocalization of VANGL2 and SCRIB, on the cell cortex. Scale bars =10 µm.*

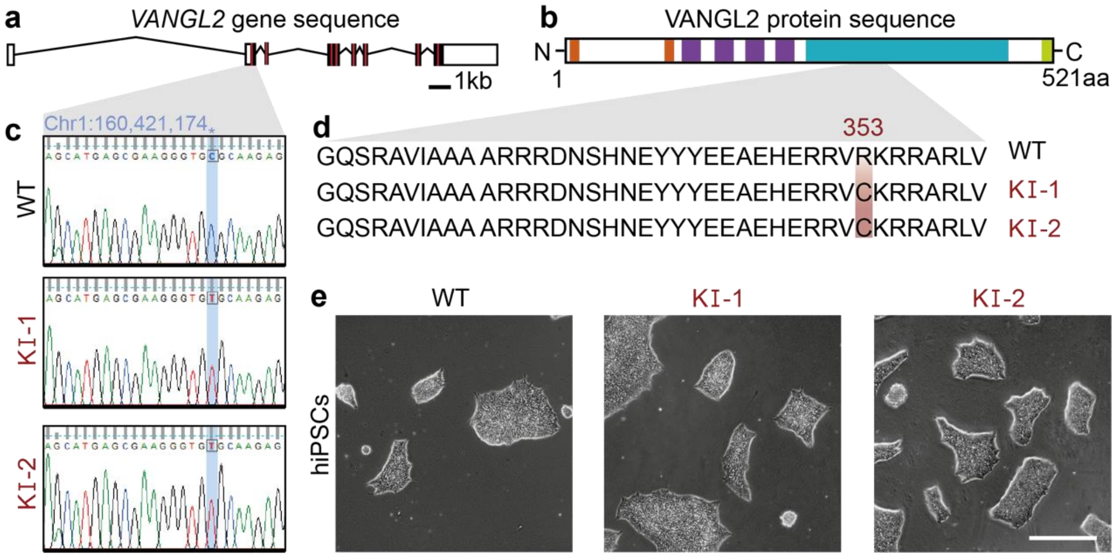

**Supplementary Figure 5: Validation of knock-in mutation in hiPSCs clones.** ***a,*** *Human VANGL2 transcript sequence, according to Ensemble database (transcript ID: ENST00000368061.3). The coding regions are coloured in black, non-coding regions in white, and the red lines indicate previously identified mutated regions.* ***b,*** *Schematic representation of the structural organisation of VANGL2, of 521 amino acids (aa). Protein domains include: two Ser/Thr phosphorylation site clusters (orange), four transmembrane domains (purple), Prickle and Dishevelled protein interaction domain (cyan), and the C-terminal PDZ binding motif (PBM) domain (green).* ***c,*** *Sanger sequencing results for WT, F1, and E2 hiPSCs clones displaying a point mutation (C>T, highlighted in blue) at Chr1:160,421,174 locus. Reads for each of the four nucleic acids, A, T, C, G are annotated in green, red, blue, and black colours, respectively.* ***d,*** *VANGL2 protein sequence (312-360 aa) for each hiPSCs clone after the introduction of the point mutation. One amino acid replacement Arginine to a Cysteine (R>C) change was found at position 353 of the protein.* ***e,*** *Brightfield images of hiPSCs colonies from WT, F1, E2 hiPSCs clones, at passage 32. Scale = 500 µm.*

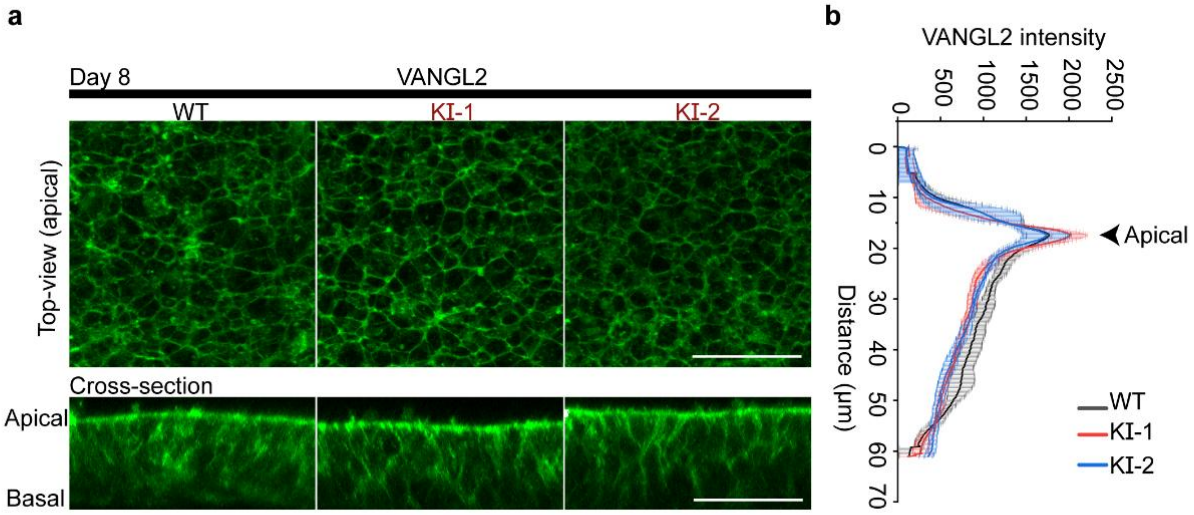

**Supplementary Figure 6: VANGL2-R353C mutation does not alter its apical localization.** ***a.*** *VANGL2 immunofluorescence showing apical and lateral views of iPSC-derived neuroepithelium from the wildtype (WT) and two knock-in (KI) lines, at day 8 of differentiation. Scale bar = 50 μm.* ***b.*** *Profile intensity quantification of VANGL2 immunofluorescence in three cultures of each line. Points represent the mean ± standard deviation. All plots are aligned so that their maximum values are in the same position.*

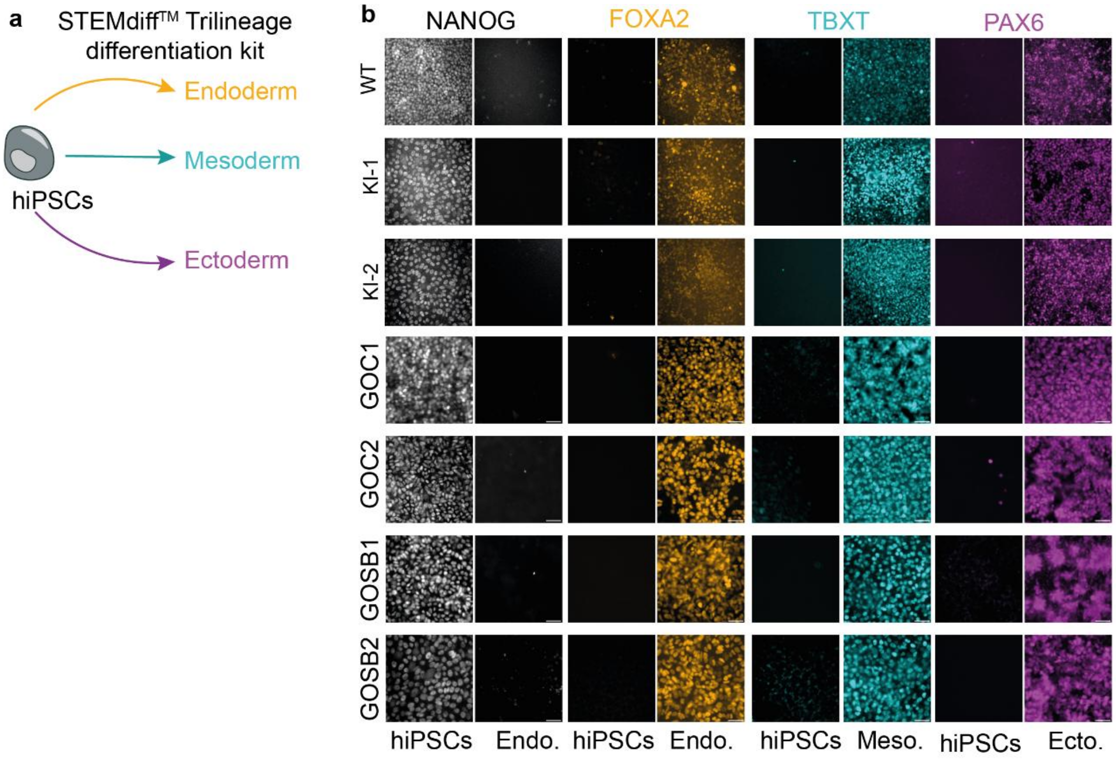

**Supplementary Figure 7: Tri-lineage differentiation of hiPSC lines.** ***a,*** *Schematic representation of tri-lineage differentiation protocol, using STEMdiff™ Trilineage Differentiation patient-derived cell lines after tri-lineage differentiation, with cells stained for NANOG (grey), FOXA2 (orange), T (cyan), and PAX6 (magenta) markers. Scale = 100 µm.*

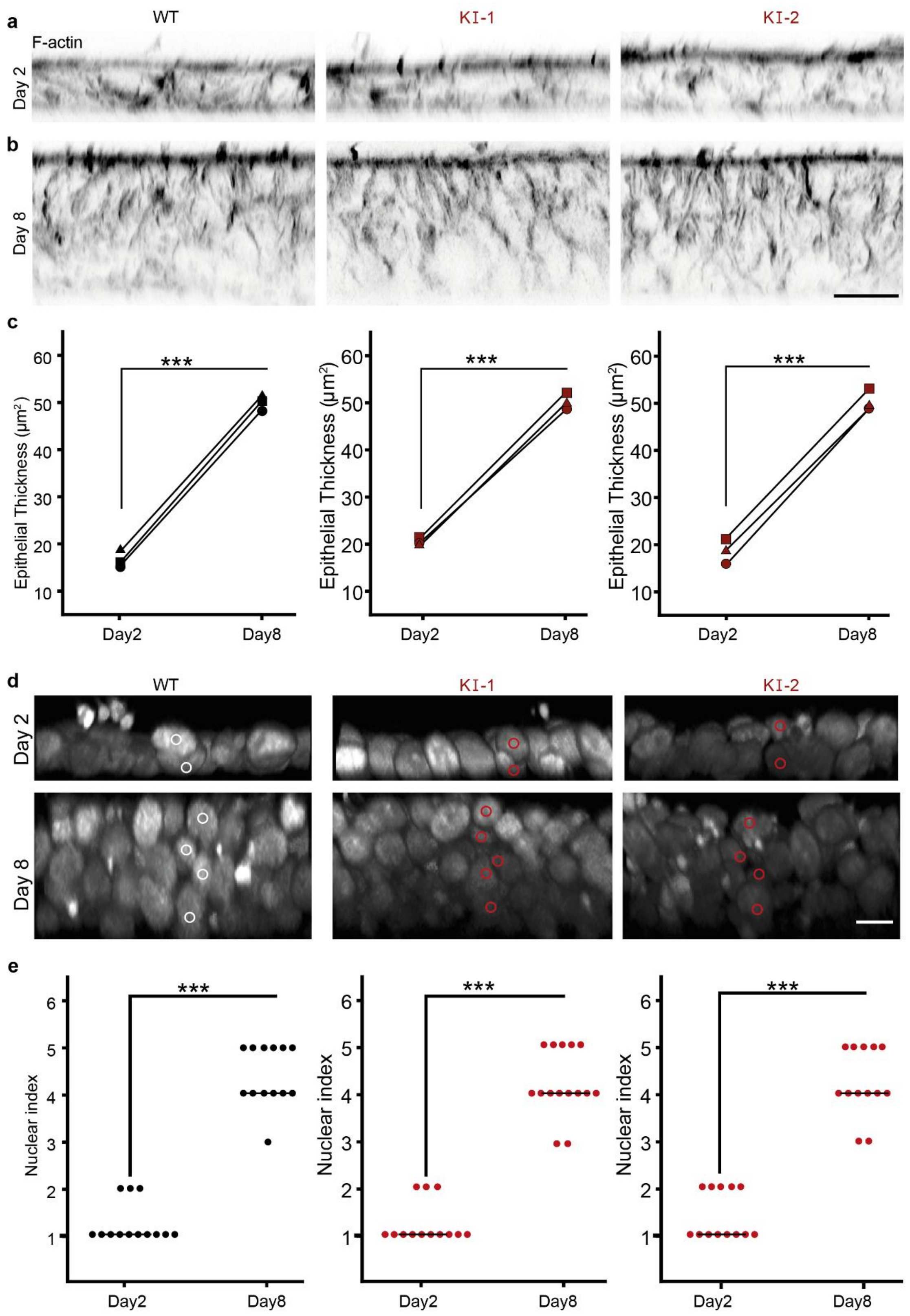

**Supplementary Figure 8: VANGL2-R353C mutation does not impair neuroepithelial apicobasal elongation.** ***a,b,*** *Optical cross-section of neuroepithelial sheets on day 2 (**a**) and*

*day 8 (**b**), after staining with F-actin (inverted grey LUT), respectively. Scale = 20 µm.* ***c,*** *Quantification of epithelial thickness on day 2 and day 8 samples, across WT and VANGL2-KI cell lines. Points represent independent plate quantification for each line. Paired T-test was performed to compare mean epithelial thickness for each cell line across day 2 and day 8 samples.* ***d.*** *Optical cross-section of neuroepithelial sheets on day 2 and day 8, after staining with DAPI (grey). Dots annotate individual nuclei across the epithelium. Scale = 20 µm.* ***e,*** *Quantification of nuclei dispersed in a row (nuclear index) of day 2 and day 8 samples. Two sample T-test performed for mean nuclear index for each cell line across day 2 and day 8 samples (***: p ≤ 0.0001).*

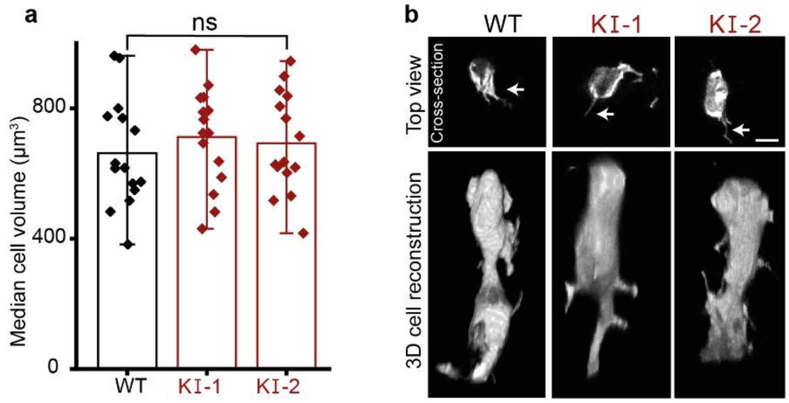

**Supplementary Figure 9: VANGL2-R353C mutation does not alter neuroepithelial cell volume.** ***a.*** *Bar graph overlaid with dot plot represent the quantification of neuroepithelial cell volume across the WT, F1, and E2 cell lines, with 15 cells analysed in each cell line. Statistical test used for individual cell volume and sub-apical protrusion length was Mann Whitney U-test with post-hoc Bonferroni (n.s.: not significant).* ***b.*** *Representative image of subapical cross-section of neuroepithelial cells, day 8 stained with Bio-Tracker (grey). Arrowheads indicate sub-apical protrusions. Scale = 20 µm.*

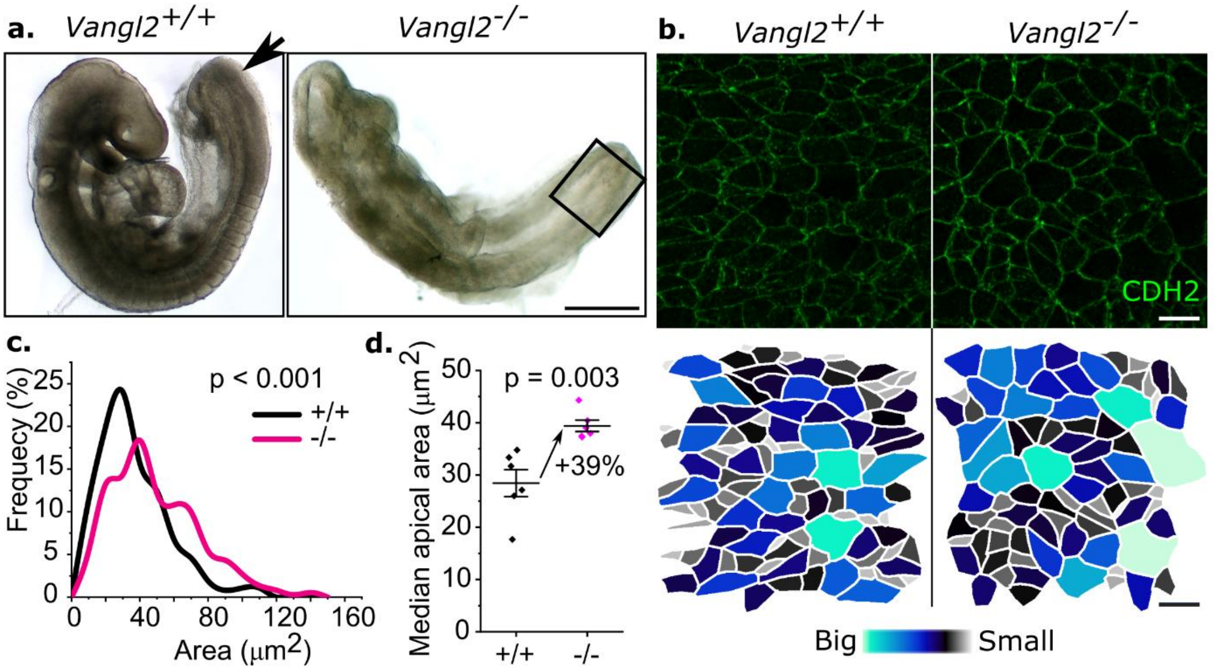

**Supplementary Figure 10: Mouse embryos globally lacking Vangl2 have larger neuroepithelial apical areas.** ***a.*** *Brightfield images of a Vangl2-knockout mouse embryo with craniorachischisis and littermate control. The arrow indicates the posterior neuropore in the control embryos, and the black box indicates the equivalent region of tissue analysed in the knockout. Scale bar = 500 µm.* ***b,*** *CDH2 (N-cadherin) immunolocalization showing the apical surfaces of neuroepithelial cells in the regions indicated in panel* ***a****, with apical areas shown as a heatmap below. Scale bars = 10 µm.* ***c,*** *Frequency plot analysis of apical areas distributions in control and Vangl2-knockout embryos. Controls = 370 cells and knockouts = 360 cells from 6 embryos per genotype, 19-23 somite stage. P value by Kolmogorov Smirnov test.* ***d,*** *Representation of the apical area data in* ***c.*** *with each point representing the median apical area of an individual embryo. P value by unpaired T-test.*

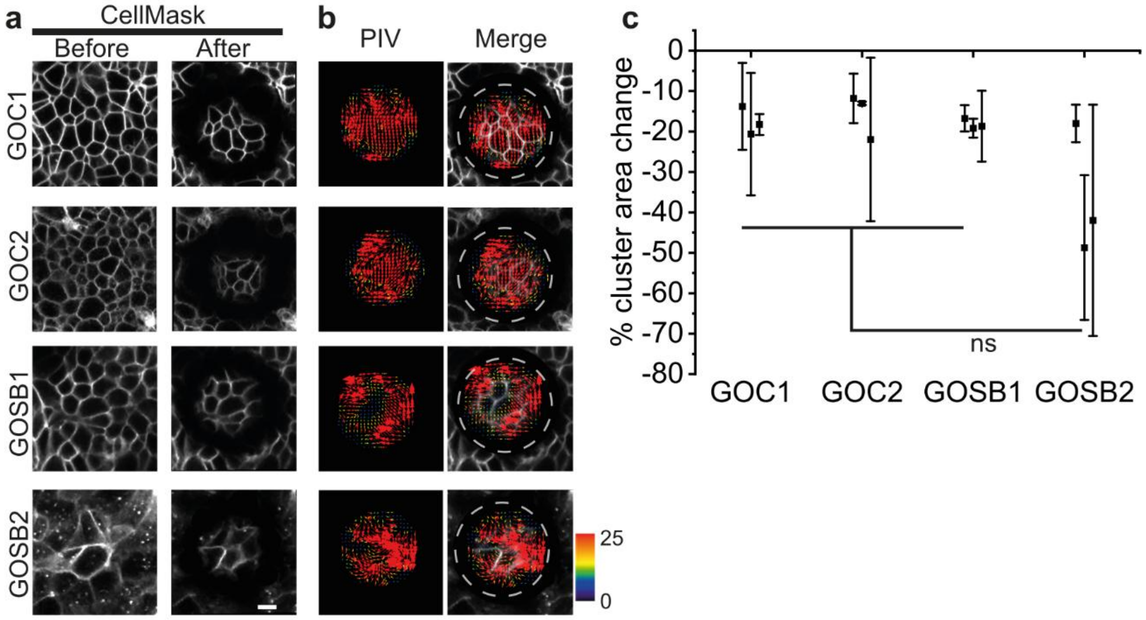

**Supplementary Figure 11: Neuroepithelial cells differentiated from the control and spina bifida patient-derived iPSC lines show equivalent apical retraction following laser ablation.** ***a,*** *Live-imaged CellMask-stained neuroepithelial sheet, day 8, before and after a linear circular laser ablation (white dash line). Scale = 20 µm.* ***b,*** *Particle image velocimetry (PIV) analysis representing the direction and magnitude of recoil.* ***c,*** *Quantification of median apical cluster constriction following laser ablation, across three independent plates per line. Points represent average recoil ± standard deviation of three ablations per plate. Mean values across were compared with one way ANOVA (n.s.: not significant).*

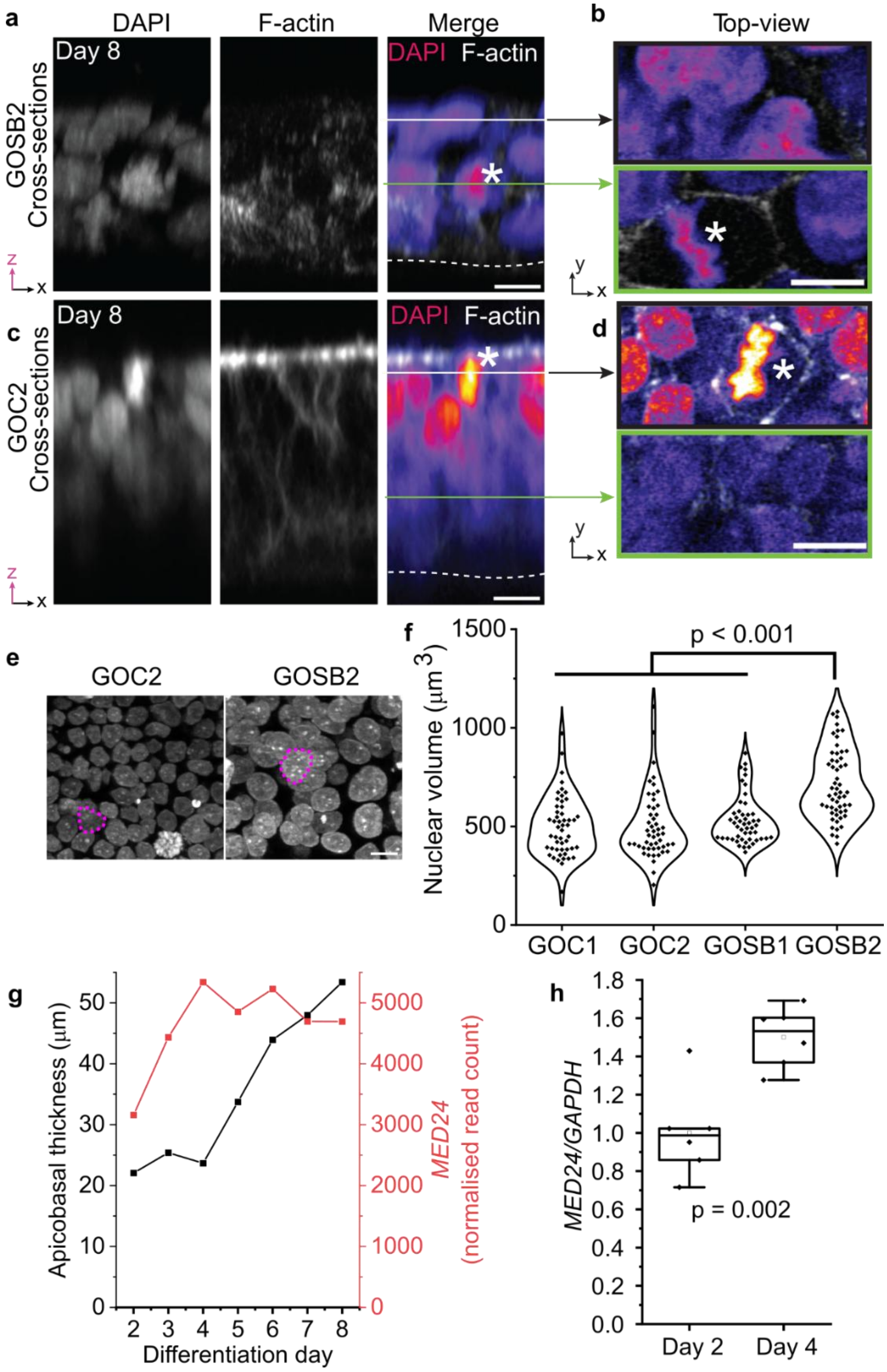

**Supplementary Figure 12: Sub-apical localisation of mitotic figure in GOSB2 spina bifida patient-derived iPSC line.** ***a, b,*** *Representative high-resolution confocal cross-section (**a**) and top-view (**b**) images of day 8 GOSB2 cultures stained against F-actin and DAPI.* ***c, d,*** *Representative high-resolution confocal cross-section (**c**) and top-view (**d**) images of day 8 GOC2 cultures stained against F-actin and DAPI. Dashed lines indicate basal site of*

*neuroepithelial culture. The arrow lines (white and green) in both merged images correspond to equivalent apical and basal slices shown in panel* **b** *and* **d***, respectively for GOSB2 and GOC2. Asterisk annotates the mitotic figure during metaphase.* ***e.*** *Visualisation of DAPI-stained nuclei on day 8 of differentiation in the indicated iPSC lines. Magenta outlines illustrate nuclear perimeter, scale bar = 10 µm.* ***f.*** *Quantification of nuclear volume on day 8 from >50 nuclei per iPSC line from one differentiation (larger GOSB3 nuclei observed in three independent differentiations). P value by ANOVA with post-hoc Bonferroni.* ***g.*** *RNASeq quantification of MED24 expression and quantification of apicobasal thickness in parallel cultures of iPSC3 cells. Note MED24 up-regulation precedes elongation.* ***h.*** *qPCR quantification of MED24 relative to GAPDH in the same iPSC3 line (6 wells in a single induction).*

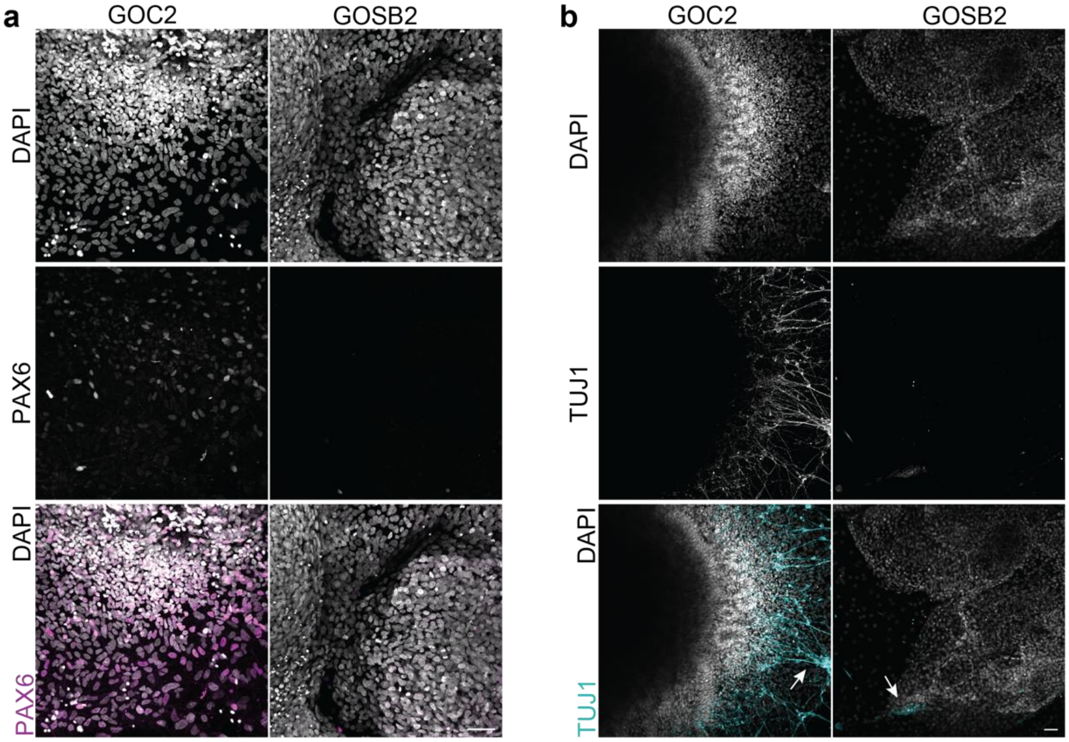

**Supplementary Figure 13: Neuronal rosette comparison between GOC2 and GOSB2 patient-derived cell lines.** ***a.*** *Representative immunofluorescence panel of day 20, neuronal rosettes stage cultures stained against PAX6 (magenta) and DAPI (grey). Scale = 50 µm.* ***b.*** *Representative immunofluorescence panel of day 20, neuronal rosettes stage cultures stained against TUJ1 (cyan) and DAPI (grey). Arrows indicate TUJ1+ outgrowths. Scale = 50 µm.*
