## Supplementary material for "Patient-specific iPSC models of neural tube defects identify deficiencies in neuroepithelial cell shape regulation and differentiation": Source data

### Slide 1
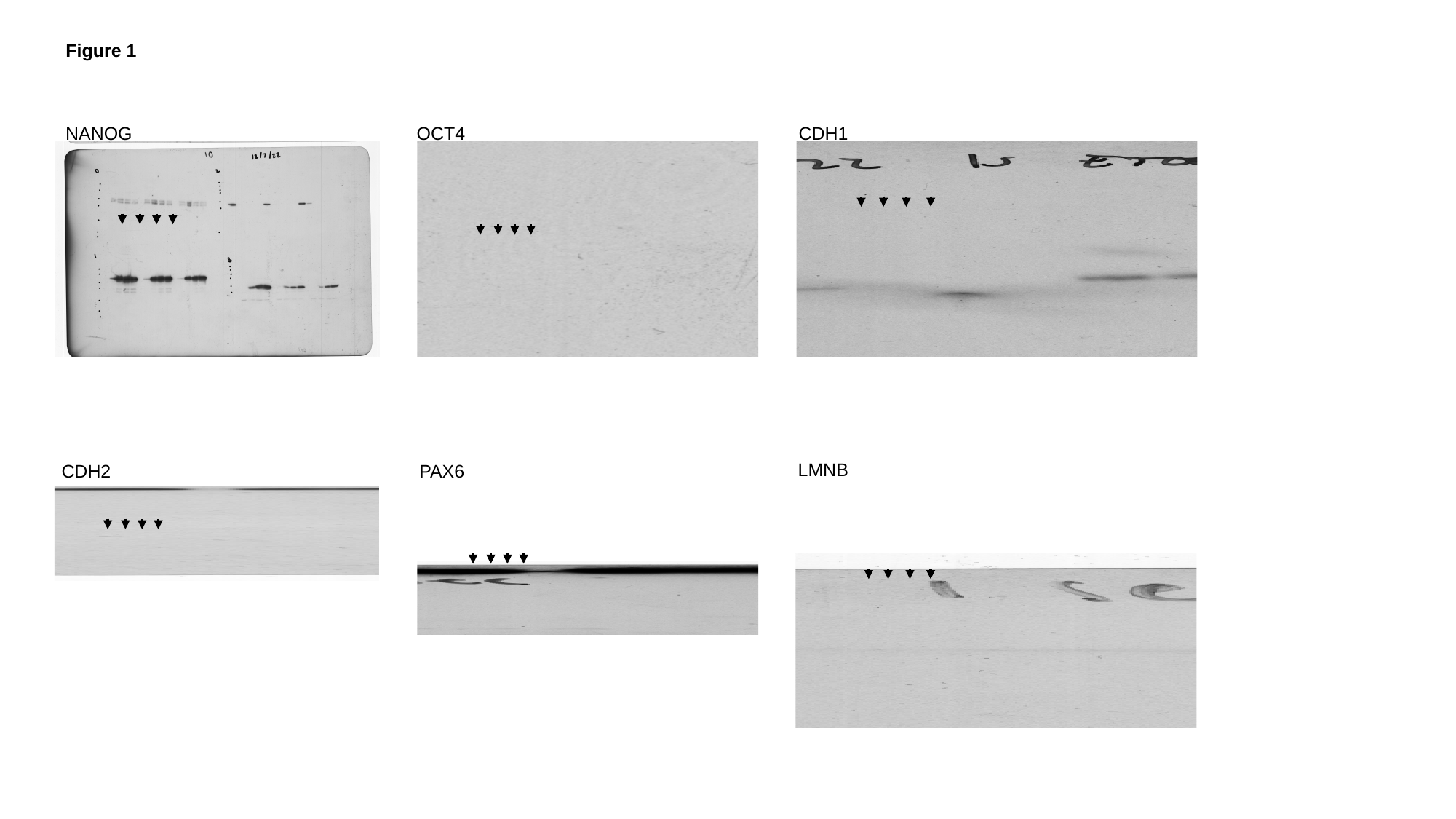

Figure 1
NANOG
OCT4
CDH1
LMNB
CDH2
PAX6

### Slide 2
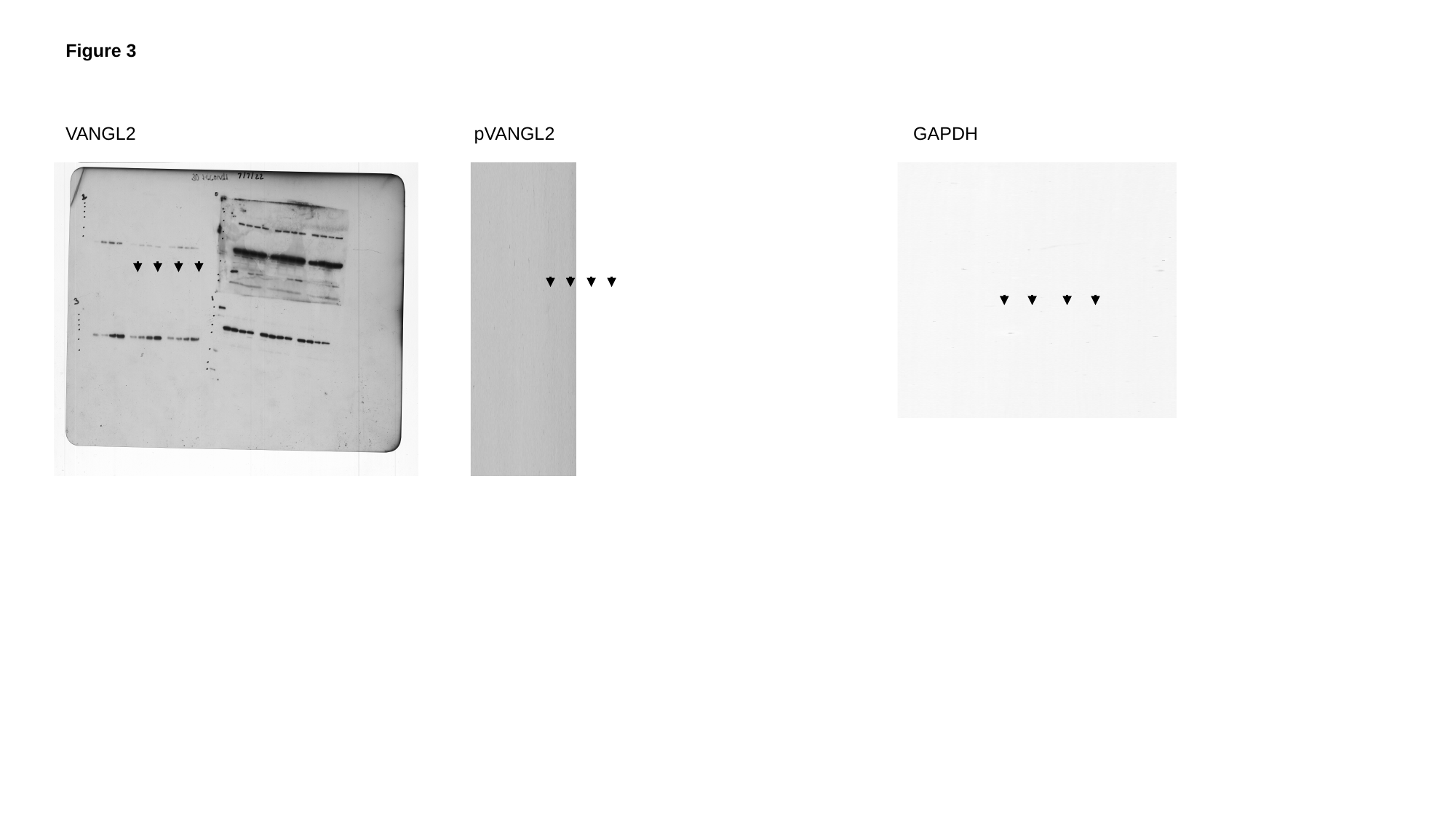

Figure 3
VANGL2
pVANGL2
GAPDH

### Slide 3
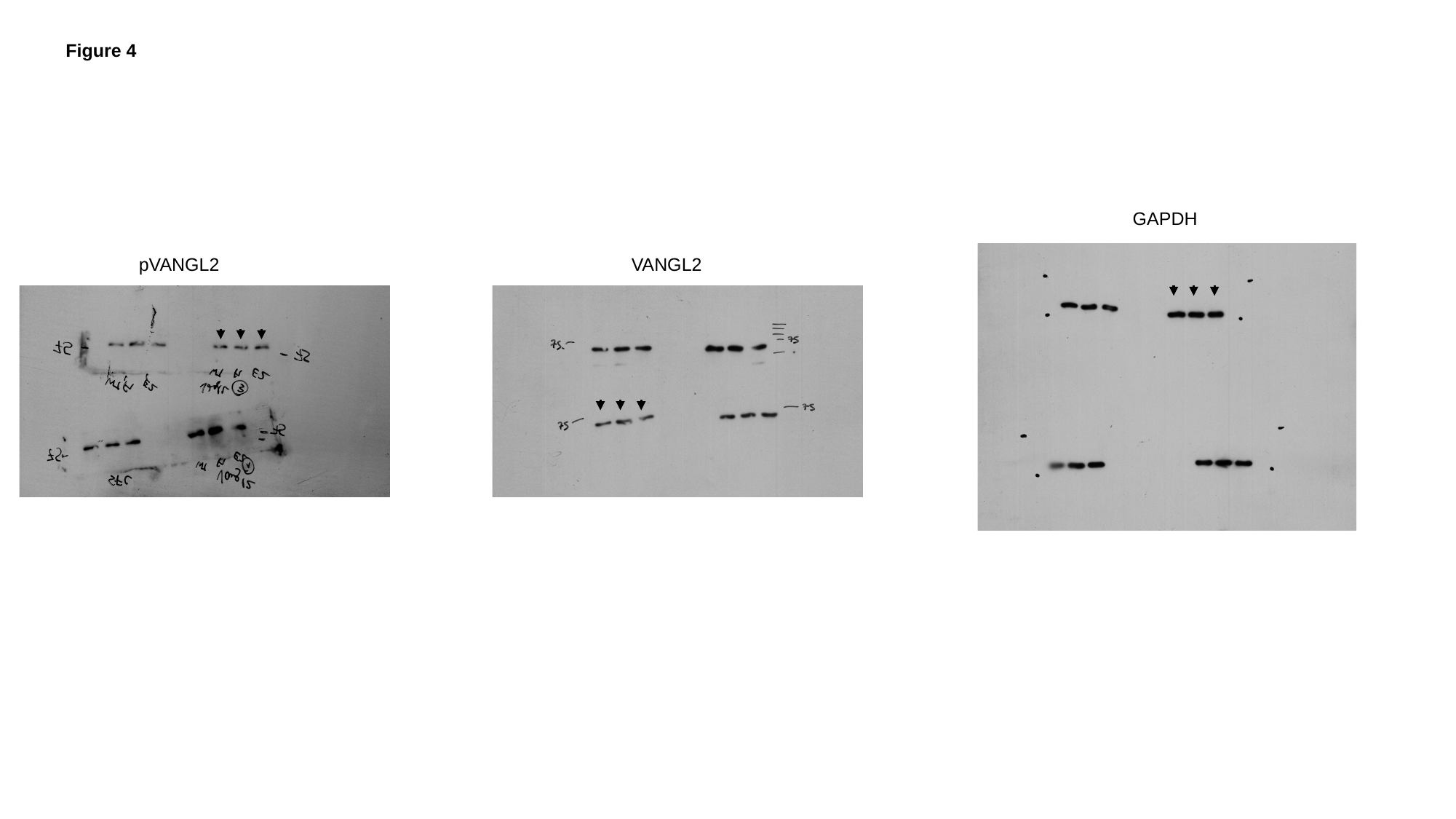

Figure 4
GAPDH
VANGL2
pVANGL2

### Slide 4
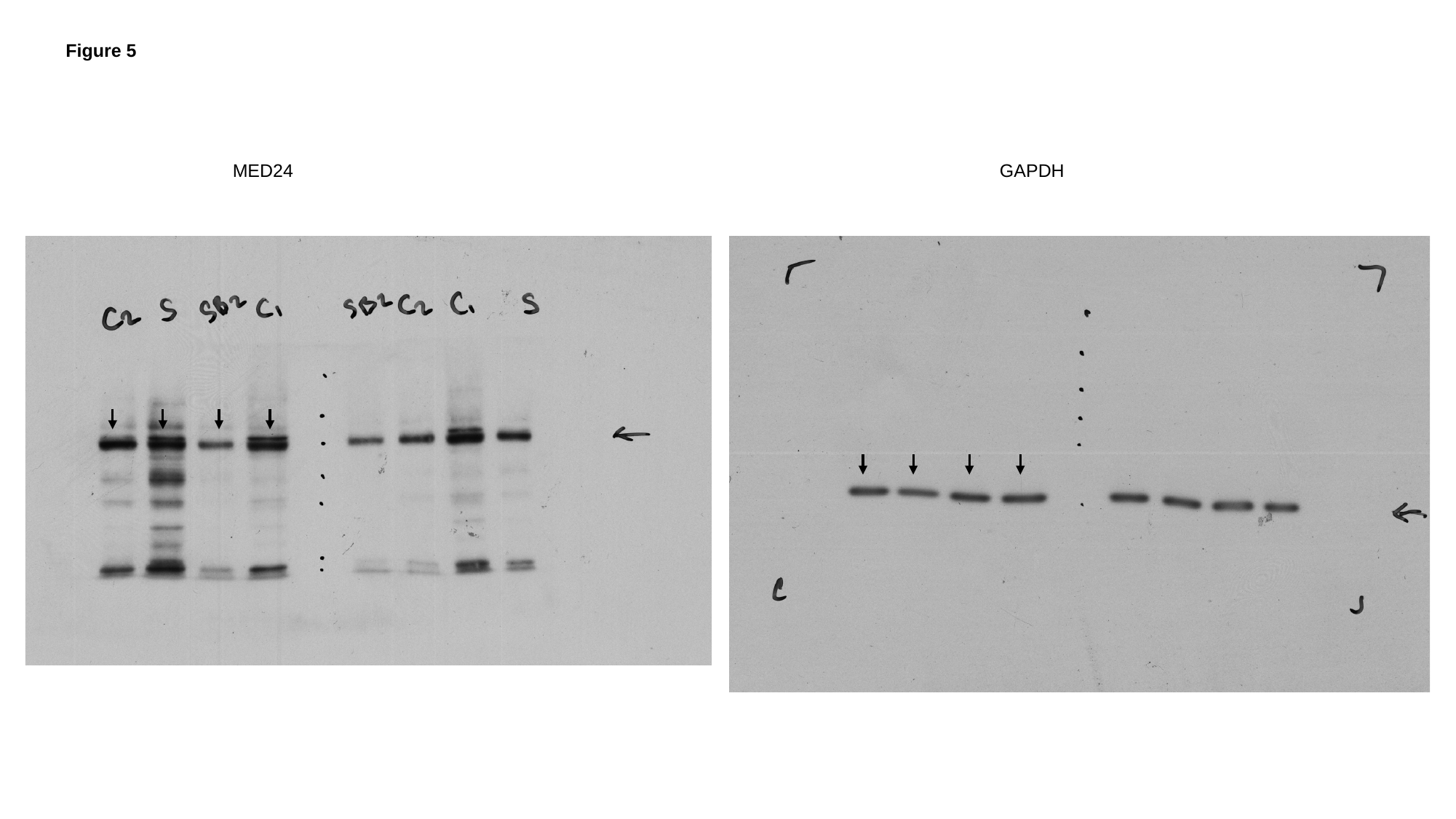

Figure 5
MED24
GAPDH
