## Appendix 1 for "Patient-specific iPSC models of neural tube defects identify deficiencies in neuroepithelial cell shape regulation and differentiation"

| **Appendix 1: Key Resources Table** | | | | |
| --- | --- | --- | --- | --- |
| **Reagent type, species or resource** | **Designation** | **Source or reference** | **Identifiers** | **Additional information** |
| gene (Homo sapiens) | VANGL2 | HGNC | HGNC:15511; NCBI Gene:57216 | R353C variant (Lei et al., 2010) knocked in; see Methods, VANGL2 knock-in cell line generation |
| gene (H. sapiens) | MED24 | HGNC | HGNC:22963; NCBI Gene:9862 | Compound heterozygous variants in GOSB2 |
| gene (Mus musculus) | Vangl2 | MGI | MGI:2135272 |  |
| genetic reagent (M. musculus) | Vangl2 knockout (global Vangl2 deletion) | doi:10.1073/pnas.2023163118 | MGI:2135272 | As previously described (Maniou et al., 2021); 19–23 somite embryos vs wild-type littermates |
| cell line (H. sapiens, female) | SFC086-03-01; iPSC2; SFC086 | doi:10.1016/j.ydbio.2022.12.002 | RRID:CVCL_RC53; hPSCreg:STBCi052-A | Fibroblast-derived iPSC (46,XX), Sendai reprogrammed; provided by Prof F. Livesey. See Table 1 |
| cell line (H. sapiens, male) | HO193b; iPSC1; GOC1 | doi:10.1038/s41563-024-01942-9 |  | Control amniocyte-derived iPSC (46,XY), microfluidic mmRNA reprogramming; from NIHR GOSH BRC. See Table 1 |
| cell line (H. sapiens, male) | HO220; GOC2 | this paper |  | Control amniocyte-derived iPSC (46,XY), microfluidic mmRNA reprogramming; NIHR GOSH BRC. See Table 1 |
| cell line (H. sapiens, female) | HO471b; GOSB1 | this paper |  | Spina bifida patient amniocyte-derived iPSC (46,XX), microfluidic mmRNA reprogramming; NIHR GOSH BRC. See Table 1 |
| cell line (H. sapiens, male) | HO541; GOSB2 | this paper |  | Spina bifida patient amniocyte-derived iPSC (46,XY), microfluidic mmRNA reprogramming; NIHR GOSH BRC. See Table 1 |
| cell line (H. sapiens, female) | 802-30F, parental; iPSC3; WT | Synthego | RRID:CVCL_E3R4; hPSCreg:CIGLi001-A | Endothelial-derived iPSC (46,XX), StemRNA 3rd Gen. Isogenic control: SpCas9-only electroporation, unedited at target locus |
| cell line (H. sapiens, female) | 802-30F, VANGL2-R353C, clone E2; KI-1 | this paper |  | Homozygous VANGL2 R353C (CGC>TGC) knock-in by CRISPR-Cas9 HDR, Synthego SO-9291367-1; see Methods |
| cell line (H. sapiens, female) | 802-30F, VANGL2-R353C, clone F1; KI-2 | this paper |  | Homozygous VANGL2 R353C (CGC>TGC) knock-in by CRISPR-Cas9 HDR, Synthego SO-9291367-1; see Methods |
| antibody | anti-ROCK1/ROCK2 (Rabbit monoclonal, clone EP786Y) | Abcam | Abcam:ab45171 | IF (1:200) |
| antibody | anti-phospho-Myosin Light Chain 2 (Ser19) (pMLC) (Rabbit polyclonal) | Cell Signaling Technology | CST:3671; RRID:AB_330248 | IF (1:200) |
| antibody | anti-SOX2 (Rabbit polyclonal) | Merck Millipore | Millipore:AB5603; RRID:AB_2286686 | IF (1:200) |
| antibody | anti-N-cadherin/CDH2 (Mouse monoclonal, clone 13A9) | Cell Signaling Technology | CST:14215 | IF (1:200) |
| antibody | anti-Lamin A/C (Mouse monoclonal, clone 636) | Santa Cruz Biotechnology | SCBT:sc-7292; RRID:AB_627875 | IF (1:200) |
| antibody | anti-Lamin B1 (LMNB) (Goat polyclonal) | Santa Cruz Biotechnology | SCBT:sc-6216; RRID:AB_648156 | IF (1:200) |
| antibody | anti-ZO-1 (Rabbit polyclonal) | Thermo Fisher Scientific | Thermo:40-2200; RRID:AB_2533456 | IF (1:200) |
| antibody | anti-Scribble/SCRIB (Goat polyclonal) | Santa Cruz Biotechnology | SCBT:sc-11049 | IF (1:200) |
| antibody | anti-VANGL2 (Rat monoclonal, clone 2G4) | Merck Millipore | Millipore:MABN750 | IF (1:200) |
| antibody | anti-PAX6 (Rabbit polyclonal) | BioLegend | BioLegend:901301; RRID:AB_2565003 | IF (1:200) |
| antibody | anti-FOXA2 (Rabbit monoclonal) | Thermo Fisher Scientific | Thermo: 701698, RRID:AB_2576439 | IF (1:200) |
| antibody | anti-TBXT (Rabbit monoclonal) | Abcam | ab209665, RRID:AB_2750925 | IF (1:200) |
| antibody | anti-β-III tubulin, TUJ1 (Mouse monoclonal, clone TUJ1) | BioLegend | BioLegend:MMS-435P; RRID:AB_2313773 | IF (1:200) |
| antibody | Donkey anti-Mouse IgG, Alexa Fluor 405 (Donkey polyclonal) | Abcam | Abcam:ab175659; RRID:AB_2687445 | IF (1:500) |
| antibody | Donkey anti-Mouse IgG, Alexa Fluor 488 (Donkey polyclonal) | Abcam | Abcam:ab150109; RRID:AB_2571721 | IF (1:500) |
| antibody | Donkey anti-Mouse IgG, Alexa Fluor 488 (Donkey polyclonal) | Abcam | Abcam:ab150109; RRID:AB_2571721 | IF (1:500) |
| antibody | Goat anti-Mouse IgG, Alexa Fluor 488 (Goat polyclonal) | Invitrogen | Invitrogen:A11029; RRID:AB_2534088 | IF (1:500) |
| antibody | Goat anti-Rabbit IgG, Alexa Fluor 488 (Goat polyclonal) | Invitrogen | Invitrogen:A11034; RRID:AB_2576217 | IF (1:500) |
| antibody | Donkey anti-Goat IgG, Alexa Fluor 488 (Donkey polyclonal) | Abcam | Abcam:ab150133 | IF (1:500) |
| antibody | Goat anti-Rat IgG, Alexa Fluor 488 (Goat polyclonal) | Invitrogen | Invitrogen:A11006; RRID:AB_2534074 | IF (1:500) |
| antibody | Goat anti-Mouse IgG, Alexa Fluor 568 (Goat polyclonal) | Invitrogen | Invitrogen:A11004; RRID:AB_2534072 | IF (1:500) |
| antibody | Goat anti-Rabbit IgG, Alexa Fluor 568 (Goat polyclonal) | Invitrogen | Invitrogen:A11011; RRID:AB_143157 | IF (1:500) |
| antibody | Donkey anti-Goat IgG, Alexa Fluor 568 (Donkey polyclonal) | Abcam | Abcam:ab175704 | IF (1:500) |
| antibody | Goat anti-Rat IgG, Alexa Fluor 568 (Goat polyclonal) | Invitrogen | Invitrogen:A11077; RRID:AB_2534121 | IF (1:500) |
| antibody | anti-phospho-VANGL2 (Ser79/Ser82/Ser84) (pVANGL2) (Rabbit monoclonal) | Sigma-Aldrich (Merck) | Sigma:SAB5701946 | WB (1:500) |
| antibody | anti-phospho-VANGL2 (Ser79/Ser82/Ser84) (pVANGL2) (Rabbit monoclonal) | Sigma-Aldrich (Merck) | Sigma:SAB5701946 | WB (1:500) |
| antibody | anti-NANOG (Mouse monoclonal, clone 23D2-3C6) | Abcam | Abcam:ab173368 | WB (1:500) |
| antibody | anti-OCT4 (Rabbit polyclonal) | Abcam | Abcam:ab19857; RRID:AB_445175 | WB (1:500) |
| antibody | anti-E-cadherin/CDH1 (Mouse monoclonal, clone 180224) | R&D Systems | R&D: MAB18381 | WB (1:500) |
| antibody | anti-GAPDH (Mouse monoclonal, clone 6C5) | Merck Millipore | Millipore:MAB374; RRID:AB_2107445 | WB (1:500) |
| antibody | anti-MED24/TRAP100 (Rabbit polyclonal) | Bethyl Laboratories | Bethyl:A301-472A | WB (1:500) |
| antibody | Goat anti-mouse immunoglobulins/HRP | Dako | Dako: P0447; RRID:AB_2617137 | WB (1:500) |
| antibody | Goat anti-rabbit immunoglobulins/HRP | Dako | Dako: P0448; RRID:AB_2617138 | WB (1:500) |
| antibody | Rabbit anti-rat immunoglobulins/HRP | Dako | Dako: P0450; | WB (1:500) |
| sequence-based reagent | VANGL2-R353C sgRNA | this paper (Synthego) | sgRNA | AUGAGCGAAGGGUGCGCAAG |
| sequence-based reagent | VANGL2-R353C HDR donor | this paper (Synthego) | ssODN donor template | CAATGAGTACTACTATGAGGAGGCTGAGCATGAGCGAAGGGTGTGCAAGAGGAGGGCCAGGTGGGTCCCTGGGGGAGAAGAGGAGAG |
| sequence-based reagent | GAPDH_F | this paper | qPCR primer | TTGGCTACAGCAACAGGGTG |
| sequence-based reagent | GAPDH_R | this paper | qPCR primer | GGGGAGATTCAGTGTGGTGG |
| sequence-based reagent | MED24_F | this paper | qPCR primer | TCTCATGGAGACAAGGACTTCA |
| sequence-based reagent | MED24_R | this paper | qPCR primer | CGCTTAGCCATAAGGTTGTTGA |
| recombinant protein | FGF2 (Animal-Free Human FGF-basic) | PeproTech | PeproTech:AF-100-18B | (20 ng/ml), neuronal differentiation |
| recombinant protein | Laminin-521 | Gibco (Thermo Fisher Scientific) | Gibco:A29249 | Substrate for final neuronal plating |
| commercial assay or kit | MycoAlert Mycoplasma Detection Kit | Lonza | Lonza:LT07-318 | Monthly mycoplasma testing |
| commercial assay or kit | RNeasy Mini Kit | Qiagen | Qiagen:74104 | RNA extraction for bulk RNA-seq |
| commercial assay or kit | RNeasy Plus Mini Kit | Qiagen | Qiagen:74134 | RNA extraction for RT-qPCR |
| commercial assay or kit | ProtoScript II First Strand cDNA Synthesis Kit | New England Biolabs | NEB:E6560L |  |
| commercial assay or kit | Luna Universal qPCR Master Mix | New England Biolabs | NEB:M3003L |  |
| commercial assay or kit | Pierce BCA Protein Assay Kit | Thermo Fisher Scientific | Thermo:23225 |  |
| commercial assay or kit | ECL Prime Western Blotting Detection Reagent | Cytiva | Cytiva:RPN2232 |  |
| commercial assay or kit | STEMdiff Trilineage Differentiation Kit | STEMCELL Technologies | STEMCELL:05230 | Tri-lineage differentiation (Supplementary Figure 7) |
| chemical compound, drug | Dorsomorphin | Tocris | Tocris:3093 | (1 µM) in neural induction medium |
| chemical compound, drug | SB431542 | Tocris | Tocris:1614 | (10 µM) in neural induction medium |
| chemical compound, drug | Y27632 (ROCK inhibitor) | Tocris | Tocris:1254 | (10 µM), 24 hours |
| software, algorithm | Fiji/ImageJ | <https://imagej.net/software/fiji/> | RRID:SCR_002285 | Image analysis |
| software, algorithm | PIV (Particle Image Velocimetry) ImageJ plugin | doi:10.1073/pnas.1106377109 |  | Tseng et al., 2012 |
| software, algorithm | ZEN Black v2.3 | Carl Zeiss Microscopy | RRID:SCR_018163 | Image acquisition |
| software, algorithm | SPSS Statistics 25 | IBM | RRID:SCR_002865 |  |
| software, algorithm | Origin 2020 | OriginLab | RRID:SCR_014212 |  |
| software, algorithm | ShinyGO 0.80 | doi:10.1093/bioinformatics/btz931 | RRID:SCR_019213 | GO enrichment analysis |
| software, algorithm | RNAfold (ViennaRNA) | doi:10.1093/bioinformatics/btl023 | RRID:SCR_024427 | RNA secondary structure prediction |
| software, algorithm | gnomAD v4.1.0 | doi:10.1038/s41586-023-06045-0 | RRID:SCR_014964 | Variant allele frequencies |
| software, algorithm | ClinVar | NCBI | RRID:SCR_006169 | NTD-associated gene lists |
| other | Matrigel Growth Factor Reduced | Corning | Corning:354277 | (0.5%) iPSC culture; (1%) neuronal differentiation |
| other | Essential 8 (E8) medium | Thermo Fisher Scientific | Thermo:A1517001 | iPSC maintenance |
| other | N2 supplement (100x) | Gibco (Thermo Fisher Scientific) | Gibco:17502-048 | Component of neural maintenance medium; see Table 2 |
| other | B27 supplement | Gibco (Thermo Fisher Scientific) | Gibco:17504-044 | Component of neural maintenance medium; see Table 2 |
| other | Dispase | Thermo Fisher Scientific | Thermo:17105041 | Passaging of neuroepithelial sheets and neural rosettes (15–20 min, 37°C) |
| other | DAPI stain | Severn Biotech | Severn Biotech:17057 | Nuclear stain |
| other | Alexa Fluor 647 Phalloidin | Thermo Fisher Scientific | Thermo:A22287 | F-actin stain |
| other | CellMask membrane dye | Invitrogen | Invitrogen:C10046 | Live membrane stain for laser ablation |
| other | CellMask Deep Red Actin Tracking Stain | Thermo Fisher Scientific | Thermo:A57245 | Live imaging (5 min in NIM) |
| other | Bulk RNA sequencing service | GENEWIZ (Azenta Life Sciences) |  | See Methods, Bulk RNA-seq |
| other | Whole exome sequencing service | Azenta Life Sciences |  | See Methods, Whole exome sequencing |
